# Whisper 2: indel-sensitive short read mapping

**DOI:** 10.1101/2019.12.18.881292

**Authors:** Sebastian Deorowicz, Adam Gudyś

**Affiliations:** Institute of Informatics, Silesian University of Technology, Gliwice, Poland

## Abstract

**Summary:** Whisper 2 is a short-read-mapping software providing superior quality of indel variant calling. Its running times place it among the fastest existing tools.

**Availability and Implementation:** https://github.com/refresh-bio/whisper

**Supplementary information:** Supplementary data are available at publisher’s Web site.

## 1 Introduction

In spite of the development of the third generation sequencing, the combination of high throughput and low error rate of short read platforms keeps them indispensable in many biological analyses (either alone, or facilitated with long reads). These are, above all, small (Kim *et al.*, 2018) and structural (Cameron *et al.*, 2019) variant calling, but also RNA-seq (Stark *et al.*, 2019), or genome assembly (Bertrand *et al.*, 2019). Since the majority of variant callers require mapping reads to the reference genome, the reliability of the latter is crucial for variant calling accuracy.

We present Whisper 2, an algorithm for short read mapping. Equipped with a novel procedure for indel handling, it offers superior accuracy in variant calling pipeline at competitive running times.

## 2 Methods

Whisper 2 follows the main idea of its predecessor (Deorowicz *et al.*, 2019)—it sorts reads which provides locality, thus cache-friendliness, when accessing a reference genome index. Mappings of individual reads found by querying the index in the main processing stage are aggregated in the postprocessing to obtain paired-end mappings. In contrast to Whisper 1, searching for long indels is not a rescue procedure performed when no close mappings for a pair was found, but is done for individual reads during sensitive major stage of the main processing. In this stage, the reference is queried with non-overlapping segments of reads to establish anchors. Reference area preceding (following) anchor position is scanned for a presence of a 7-symbol prefix (suffix) of the read with one mismatch allowed. Among all candidates, the one maximizing the affine score is selected. The reference span to be scanned, thus the maximum length of detected indels, is by default 50. More details are given in Supplementary Section 1.

## 3 Results

Performance evaluation of mapping algorithms was done in a variant calling pipeline. Investigated packages, i.e., BWA-MEM (Li, 2013), Minimap 2 (Li, 2018), Whisper 1 (Deorowicz *et al.*, 2019), and Whisper 2 were combined with Strelka 2 variant caller, which offers superior accuracy and execution times (Kim *et al.*, 2018). Additionally, we examined Graph Genome Pipeline (GGP, Rakocevic *et al.* (2019)), which was reported to offer excellent variant calling sensitivity by mapping reads against a genome graph. Strelka 2 was configured to detect indels of length up to 100 instead of default 49—this setting rendered better results for all investigated mappers (Supplementary Figures 25–27). Benchmarking was done with a use of samples HG001 and HG005 from Genome in a Bottle Consortium (GiaB; Zook *et al.* (2019)) and SynDip (Li *et al.*, 2018) with both, GRCh37 and GRCh38 references.

Figure 1 shows the results of variant calling. When analyzing SNPs, one can see, that for given reference all algorithms offered similar precision and differed mostly in recall. The best results were reported for GGP, then came BWA and Whisper 2 (ex-equo) which were followed by Whisper 1 and Minimap 2. In indels, Whisper 2, BWA, and Minimap 2 rendered almost the same results, being superior in terms of precision in all benchmarks. In recall, they fell between GGP (best) and Whipser 1 (worst) on GiaB samples, while in SynDip they left both competitors behind. An important observation is that recall values for GiaB samples were noticeably greater than those of SynDip. This was parially due to fact, that in the former, indels are no longer than 50 bp, while the latter contained variants with even hundreds of nucleotides, which were more challenging to call.

**Fig. 1.**
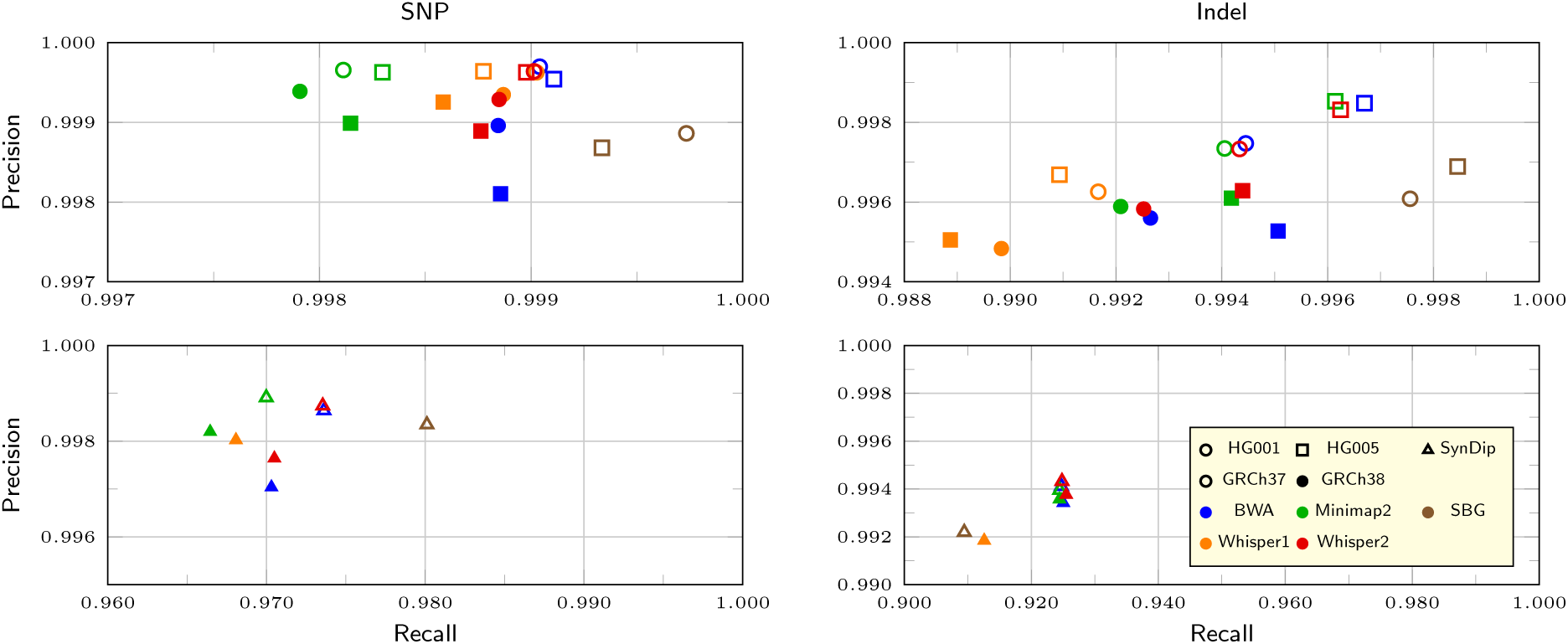
Results of variant calling presented as recall versus precision graphs. Graph Genome Pipeline supports GRCh37 reference only; Whisper 1 failed to execute on SynDip with GRCh37

In Figure 2-left, one can see the performance of calling SynDip indels of a given length *L*. For short deletions (*L* ≤ 50) Whisper 2, BWA, and Minimap 2 performed very similarly and were consistently better than GGP. In contrast, for short insertions, GGP was the leader, with Whisper 2 coming right behind it. For longer indels (51 ≤ *L* ≤ 100), the variation of F1 scores became larger. Though, one can see that Whisper 2 was the leader in deletions and held second place after GGP in insertions.

**Fig. 2.**
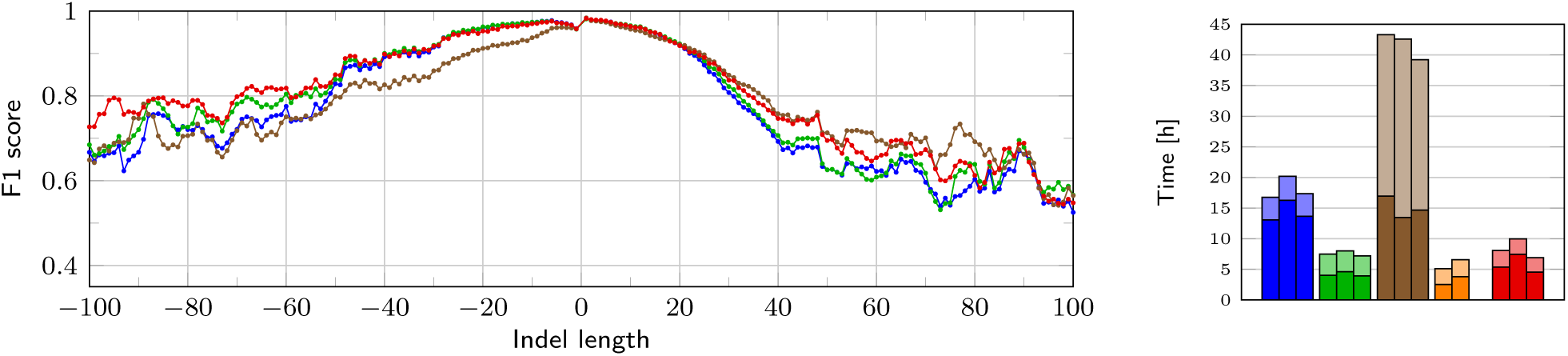
**Left:** Five-sample moving average of F1 scores for SynDyp insertions (positive) and deletions (negative) with GRCh37 reference. **Right:** Execution times at 12-core CPU for, respectively, HG001, HG005, and SynDip with GRCh37. Solid bars represent mapping, transparent include SAM to BAM conversion (if necessary), BAM sorting, and variant calling

The comparison of execution times is presented in Figure 2-right. Minimap 2 was the fastest algorithm that processed all benchmarks. However, Whisper 2 was only slightly inferior. Both of them allowed calling variants from reads in below 10 hours (at a modern workstation), while BWA-based pipeline required twice the time. Graph Genome Pipeline was the slowest with analysis time of more than 40 hours per benchmark.

## 4 Conclusions

The new method for handling indels in Whisper 2 resulted in improved variant calling performance. Presented software was on par with competitors in terms of precision and recall, revealing its potential for longer (51 ≤ *L* ≤ 100) insertions and deletions. In the former, it was second best after several times slower Genomic Graph Pipeline, in the latter it was the leader. Execution times of Whisper 2 were comparable to Minimap 2—the fastest among analyzed algorithms.

## Supporting information

Supplementary Section

