## Supplementary Section for "Whisper 2: indel-sensitive short read mapping"

### Supplementary material for article: Whisper 2: indel-sensitive short read mapping

Sebastian Deorowicz  Adam Gudyś

December 18, 2019

**2 Examined programs** **4**

**3 Datasets** **6**

**4 Environment** **7**

**5 Additional results** **8**

### 1 Methods

#### 1.1 Introduction

At the beginning let's take a brief overview of Whisper 1 algorithm.

In the first phase (preprocessing), reads are distributed into hundreds of bins according to their prefixes.

The second phase (main processing) consists of several major stages. In the $i$th major stage, the mappings to a reference genome with at most $i$ errors (mismatches or indels) are determined. It is guaranteed that such mappings, if exist, will be found. The reads with known mappings are not passed to the next major stage. The number of major stages is upper-bounded by the maximal allowed number of errors, $k$, (given as the program parameter, or determined automatically according to the read lengths). After this, the reads without known mappings fall into an additional *sensitive* major stage, in which the mappings with up to $ck$ errors (by default $c = 2.5$) are looked for, but now without guarantee that all existing such mappings will be found. The $i$th major stage is composed of $(i + 1)$ minor stages. The *sensitive* major stage is composed of $(k + 1)$ minor stages. In each minor stage, we assume that some segment of the read must match exactly some genome fragment. The fixed segments in all minor stages of a given major stage are not overlapping.

In the final stage (postprocessing), the mappings of individual reads from a pair are aggregated to form paired-end mappings. Moreover, if for one read of a pair we do not know the mapping (or if the mappings of the paired reads are distant), the paired mappings are determined using various techniques, including: ($i$) scanning the region close to one read mapping for the mapping (with more errors than $k$) of the other read; ($ii$) scanning this region with clipping of the other read.

#### 1.2 Looking for indel in the main stage

The main novelty of Whisper 2 is in the *sensitive* major stage. We implemented aggressive looking for mapping of the read with 1 or 2 (possibly long) indels. Below we describe the algorithm using an example.

Let us assume that the current read $r = r_0 r_1 \ldots r_{\ell-1}$ is of length $\ell$ and that in the current minor stage we know all exact mappings of read segment $r_p r_{p+1} \ldots r_{p+s-1}$ in the genome ($s$ is segment size). Let one of such mapping be $\mathcal{G}_q \mathcal{G}_{q+1} \ldots \mathcal{G}_{q+s-1}$. Let the maximum allowed indel length be $m$. Just for simplicity of presentation we will also assume that $m < p$ but the algorithm handles also the other case.

Here we show how we try to find the mapping of the left part (prior to $p$ position) of the read. The first step is a determination of a candidate list $C$ of mappings of a short read prefix, i.e., $e = r_0 r_1 \ldots r_6$ (the length of the prefix is an algorithm parameter) with at most one mismatch. We verify here every 7-symbol long substring of genome fragment $\mathcal{G}_{p-m} \ldots \mathcal{G}_{p+m}$.

Then, we evaluate the candidates as follows. Let the current candidate mapping of the read prefix is $t$, which means that $e$ matches $\mathcal{G}_t \mathcal{G}_{t+1} \ldots \mathcal{G}_{t+6}$ with at most one mismatch. For this candidate mapping the deletion length will be $d = q - t - p$ (negative deletion length means insertion). Now we construct two vectors representing the number of mismatches starting from both parts of the processed read fragment. The vector $M_0$ is defined such that $M_0[x]$ is the number of mismatches between $r_0 \ldots r_{x-1}$ and $\mathcal{G}_t \ldots \mathcal{G}_{t+x-1}$. The vector $M_1$ is defined similarly: $M_1[x]$ is the number of mismatches between $r_{p-x} \ldots r_{p-1}$ and $\mathcal{G}_{q-x} \ldots \mathcal{G}_{q-1}$. Knowing the current deletion length $d$ and both $M_*$ vectors we can easily find the position of the indel that maximizes the matching score. After inspecting all candidates from $C$ we pick the one that maximizes the affine score of mapping of the left read part. For completeness we calculate also the cost of mapping the left part with just mismatches.

We proceed in the same way for the right read part, i.e., starting from $r_{p+s}$. The best mappings of the left and right parts are then combined into a mapping of $r$ and the affine score is calculated. This score is compared with the minimal allowed score in the *sensitive* major stage, which is the mapping with $ck$ mismatches and if it is not smaller, the mapping is preserved for handling in the postprocessing phase.

A drawback of the above-described aggressive looking for indels is its time consumption. Nevertheless, the gain in the read mapping quality is significant, which is shown in the evaluation of the variant calling pipelines.

#### 1.3 Other improvements

Many small improvements were also implemented in the new version. Here we enumerate just a few of them.

First, we implemented the "polishing" of mappings containing a pair of close and very short indels. Such artifacts are results of our simplified mapping algorithm (not supporting affine score) implemented in the main processing phase. Polishing them, improves the performance of variant callers, both in terms of running time and quality.

Second, we implemented an improved evaluation of the mappings found during the main processing phase. In Whisper 1, the mappings in the postprocessing stage are compared using the linear score. In Whisper 2, the mappings (except for the simplest cases, in which we have perfect mappings) are reevaluated using affine score before further processing.

Third, we significantly sped up the mapping of reads of lengths in range $[128, 255]$ when the CPU does not support the AVX2 vector extensions.

### 2 Examined programs

The following programs were used in the experimental part. The running parameters are also given.

- BWA-MEM v. 0.7.15-r1140

```
bwa mem -t 12 <ref_genome> <r1.fastq> <r2.fastq> > <output.sam>
```

  The pairs of files were mapped pair by pair.

- GGP

  The Graph Genome Pipeline tools have been downloaded from https://www.sevenbridges.com/graph-genome-academic-release/ as Docker files. The mapping (version 0.9.1) and variant calling (version 0.5.20) was performed as suggested by the documentation:

```
/usr/local/bin/aligner --vcf SBG.Graph.B37.V6.rc6.vcf.gz
   --reference GRCh37_decoy.fasta -q <input_1.fastq.gz> -Q <input_2.fastq.gz>
   -o <sample.bam> --read_group_sample 'SAMPLE_READ_GROUP'
   --read_group_library 'lib' --threads 12
/usr/local/bin/reassembly_variant_caller -b <sample.sort.bam>
   -f GRCh37_decoy.fasta -g SBG.Graph.B37.V6.rc6.vcf.gz -v <results.vcf>
```

- Minimap2 v. 2.17-r941

```
minimap -t 12 -p -1 <r1.fastq.gz> -2 <r2.fastq.gz> -o <output.sam>
   -I <ref_genome>
```

  The pairs of files were mapped pair by pair.

- Sambamba v. 0.7.0

```
../sambamba/sambamba sort -t 12 -l 3 -m 64GB -o <sorted_bam_file>
   <in_bam_file>
```

  The BAM files are sorted (and converted from SAM, if necessary) using Sambamba.

- Strelka v. 2.10

  Strelka 2 was used with default options except for max. indel size set to 100 in some experiments.

- Whisper v. 1.0

```
whisper -t 12 -store-BAM -gzipped-SAM 3 -out <output.sam> <ref_genome>
   @<reads.txt>
```

  The file <reads.txt> contains pairs of names of files to be mapped separated by a semicolon, e.g., in case of 3 pairs they are processed in a single run. Sample contents of <reads.txt>:

```
r1_1.fastq.gz;r1_2.fastq.gz
r2_1.fastq.gz;r2_2.fastq.gz
```

  The additional parameters were adjusted to read lengths:

  - -e 7 -sens-factor 2.5 for HG005 dataset
  - -e 5 -sens-factor 2.5 for HG001 and SynDip datasets

- Whisper v. 2.0
  `whisper -t 12 -store-BAM -gzipped-SAM 3 -out <output.sam> <ref_genome> @<reads.txt>`
  The file `<reads.txt>` contains pairs of names of files to be mapped separated by a semicolon, e.g., in case of 3 pairs they are processed in a single run. Sample contents of `<reads.txt>`:
  ```
  r1_1.fastq.gz;r1_2.fastq.gz
  r2_1.fastq.gz;r2_2.fastq.gz
  ```

  The parameter `-max-indel-size 100` was used in some experiments. In general Whisper 2 should be (and was) used with default max. indel size, when the following variant calling tool determines only indels up to 50 bp. Nevertheless, when the variant calling tool is able to determine longer indels, setting `-max-indel-size 100` in Whisper 2 results in slightly better results.

### 3 Datasets

##### Reference genomes

Reference genomes with decoys were downloaded from:

- GRCh37: ftp://ftp-trace.ncbi.nih.gov/1000genomes/ftp/technical/reference/phase2_reference_assembly_sequence/hs37d5.fa.gz
- GRCh38: ftp://ftp.ncbi.nlm.nih.gov/genomes/all/GCA/000/001/405/GCA_000001405.15_GRCh38/seqs_for_alignment_pipelines.ucsc_ids/GCA_000001405.15_GRCh38_no_alt_analysis_set.fna.gz

### HG001

The dataset was downloaded from PrecisionFDA Truth Challenge (https://precision.fda.gov/challenges/truth):

- HG001-NA12878-50x_1.fastq.gz
- HG001-NA12878-50x_2.fastq.gz

The reads are of length 2x148 bp. The approx. coverage is 50.

### HG005

The dataset was downloaded from Genome in a Bottle Project:

- ftp://ftptrace.ncbi.nlm.nih.gov/giab/ftp/data/ChineseTrio/HG005_NA24631_son/HG005_NA24631_son_HiSeq_300x/basespace_250bps_fastqs/150424_HG005_Homogeneity_02_FCA-22108087/

The reads are of length 2x250 bp. The approx. coverage is 50.

##### SynDip

The dataset was downloaded from the European Nucleotide Archive (accession PRJEB13208):

- ftp://ftp.sra.ebi.ac.uk/vol1/fastq/ERR134/006/ERR1341796/ERR1341796_1.fastq.gz
- ftp://ftp.sra.ebi.ac.uk/vol1/fastq/ERR134/006/ERR1341796/ERR1341796_2.fastq.gz

The truth dataset was taken from https://github.com/lh3/CHM-eval/releases.

### 4 Environment

The computer used in test were of the following configuration:

- 2 Intel Xeon E5-2670 v3 CPU, 12 cores per CPU, each clocked at 2.3 GHz,
- 256 GiB RAM,
- 1 HGST Ultrastar HC510 HDD of size 10 TB, hdparm -t reported read speed 238 MB/s.

### 5 Additional results

#### 5.1 SynDip evaluation for default maximal indel length in Strelka

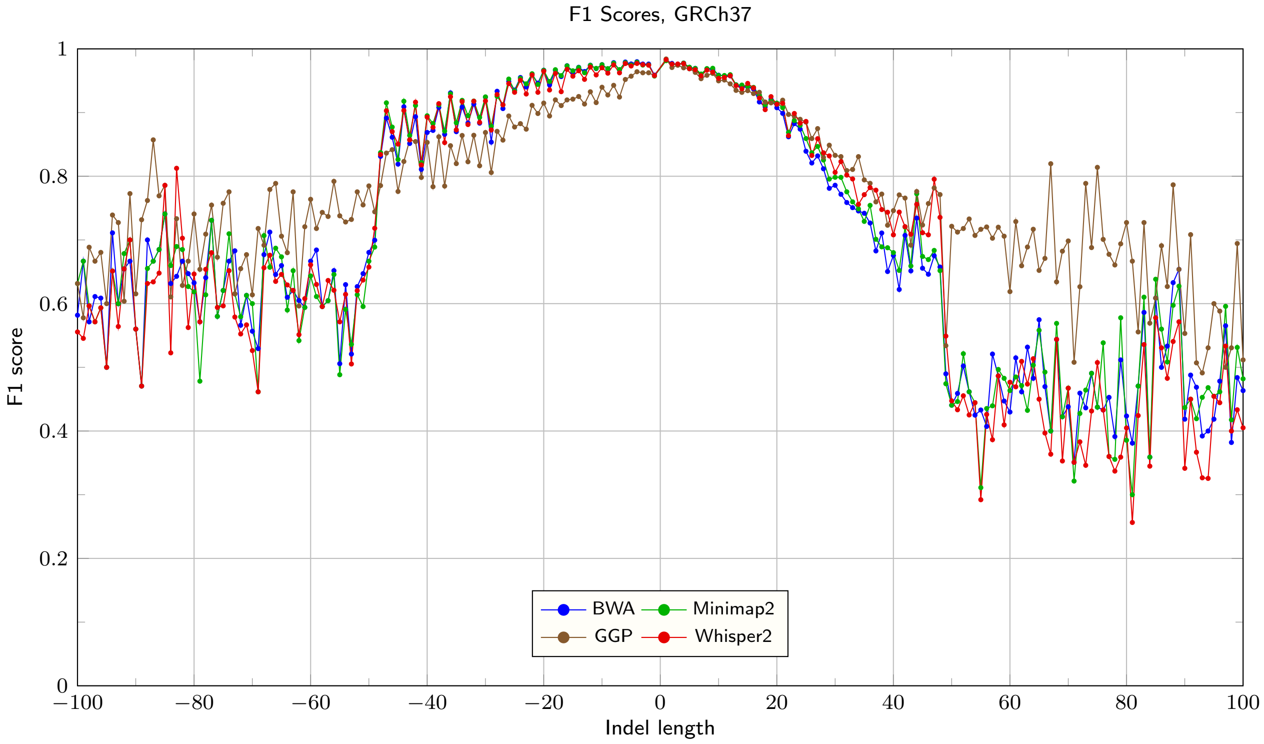

Figure 1: F1 Scores for GRCh37, Strelka max. indel length set to 49 (default)

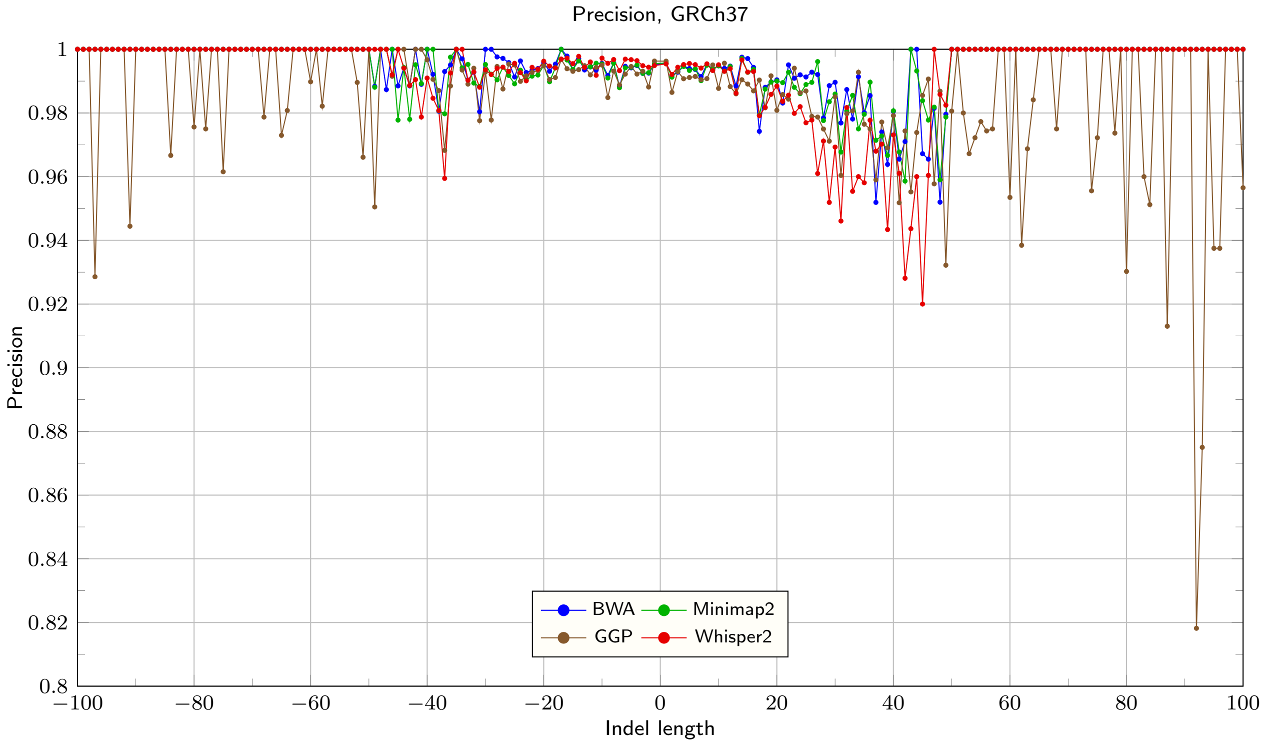

Figure 2: Precision for GRCh37, Strelka max. indel length set to 49 (default)

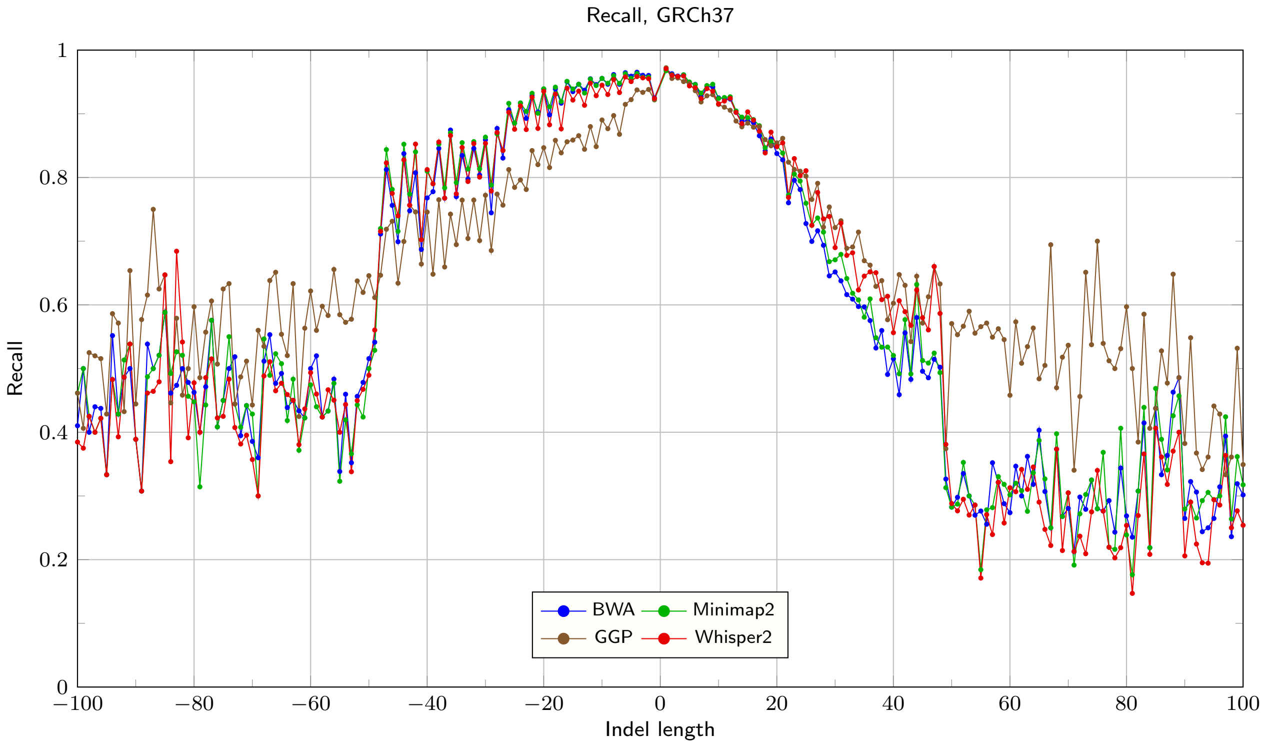

Figure 3: Recall for GRCh37, Strelka max. indel length set to 49 (default)

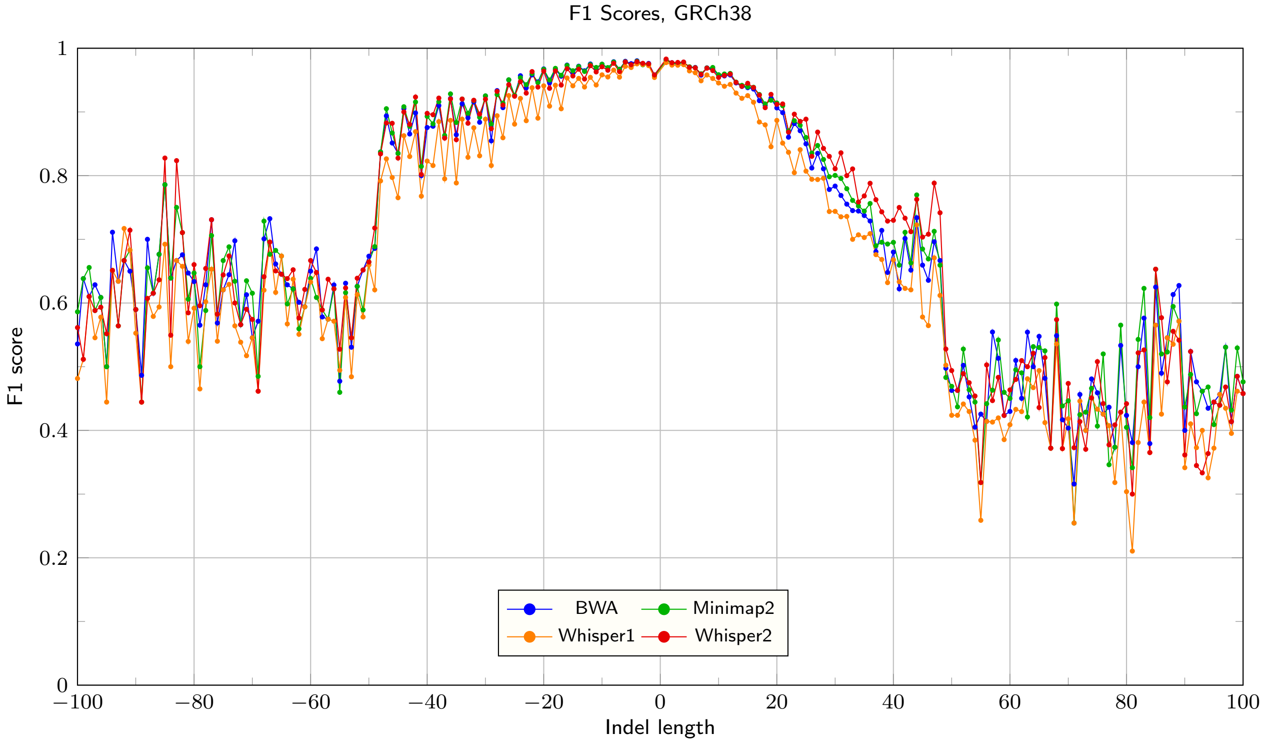

Figure 4: F1 Scores for GRCh38, Strelka max. indel length set to 49 (default)

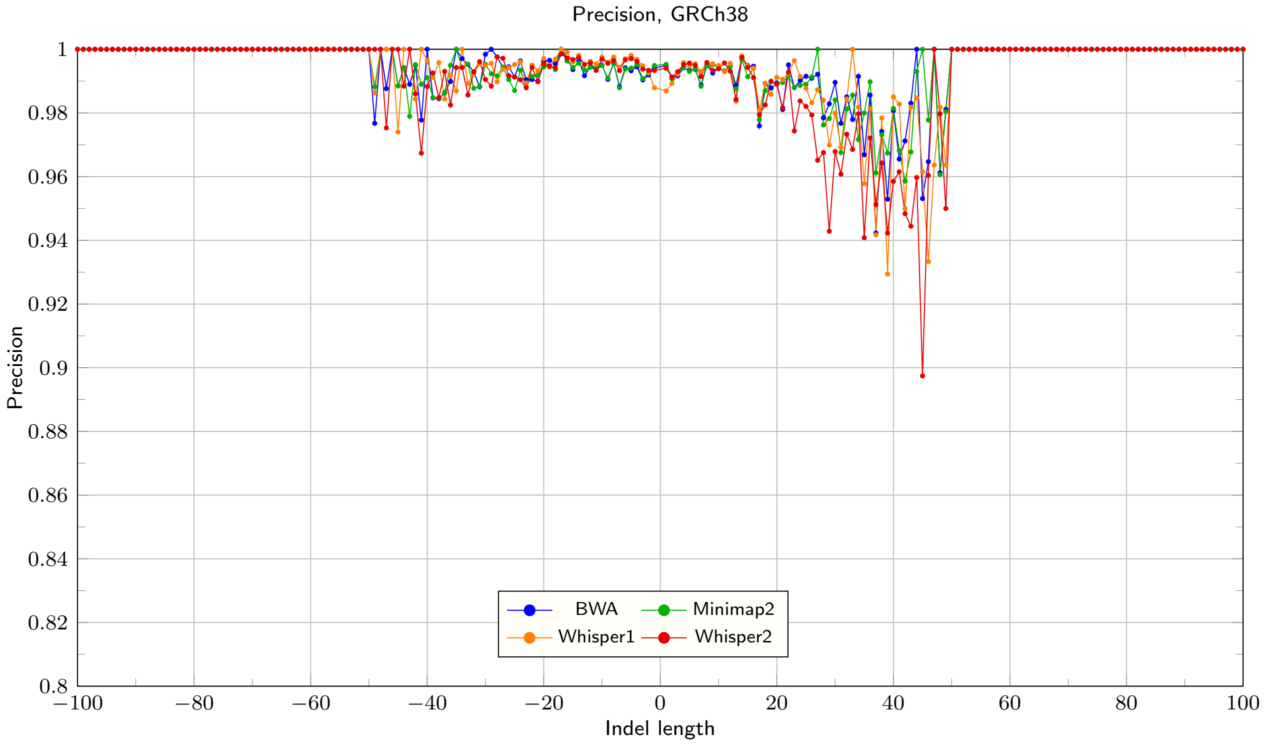

Figure 5: Precision for GRCh38, Strelka max. indel length set to 49 (default)

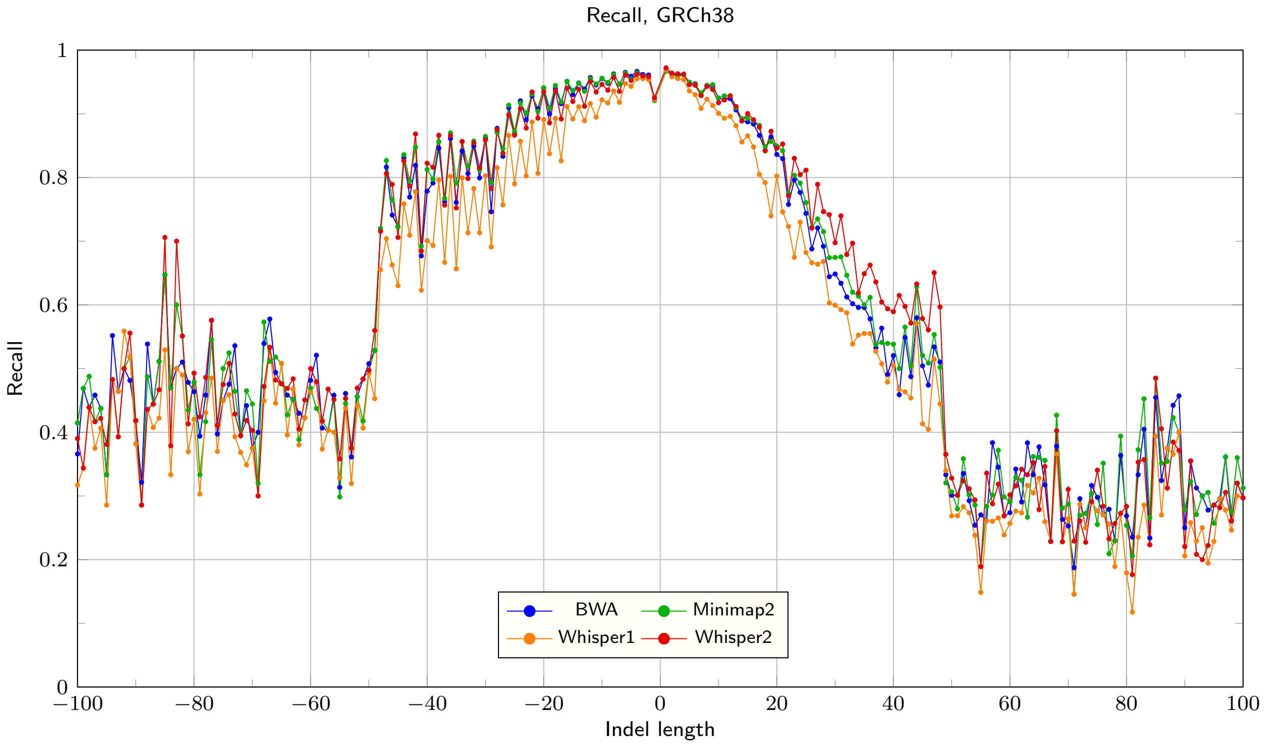

Figure 6: Recall for GRCh38, Strelka max. indel length set to 49 (default)

#### 5.2 SynDip evaluation for expanded maximal indel length in Strelka

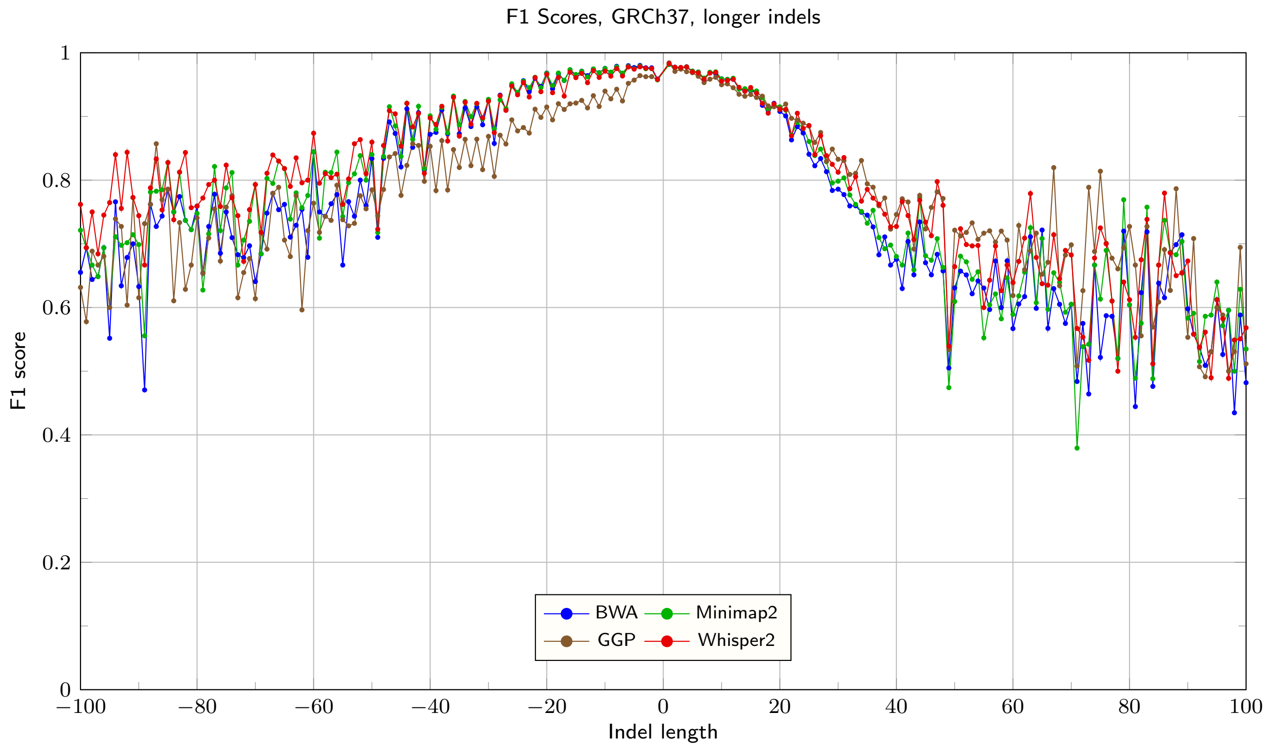

Figure 7: F1 Scores for GRCh37, Strelka max. indel length set to 100

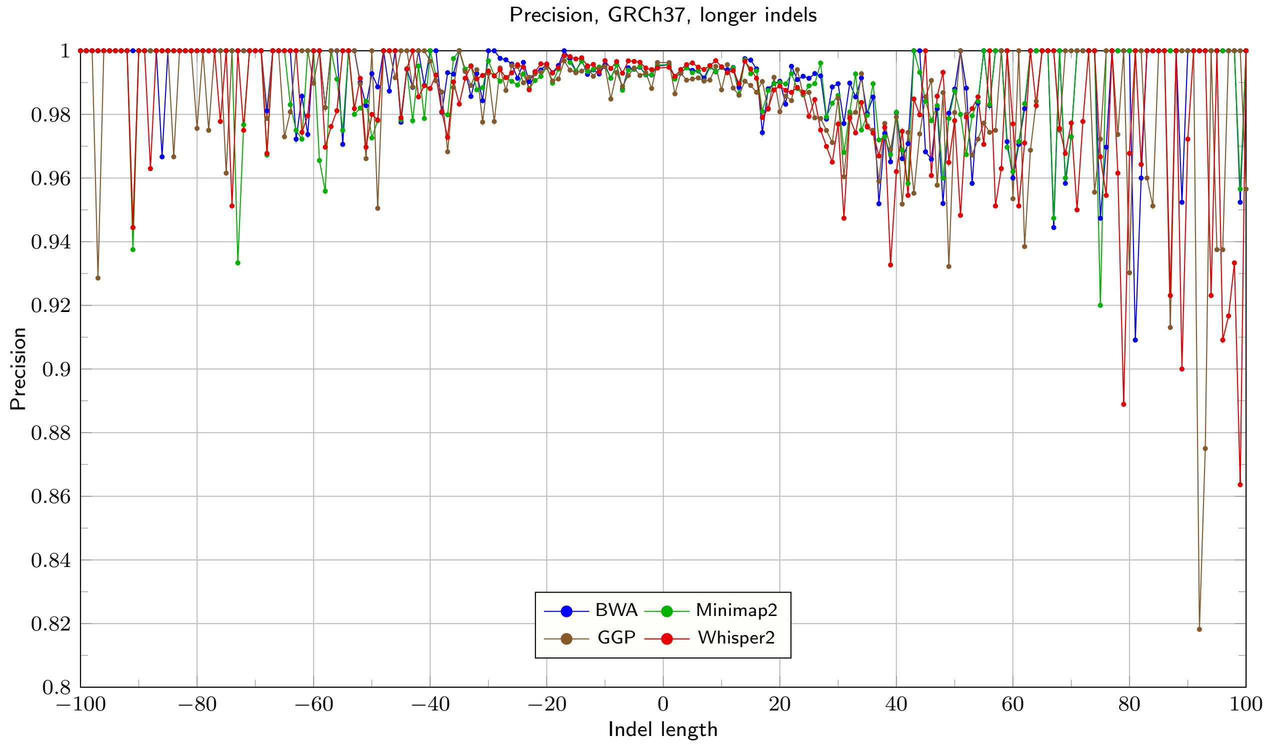

Figure 8: Precision for GRCh37, Strelka max. indel length set to 100

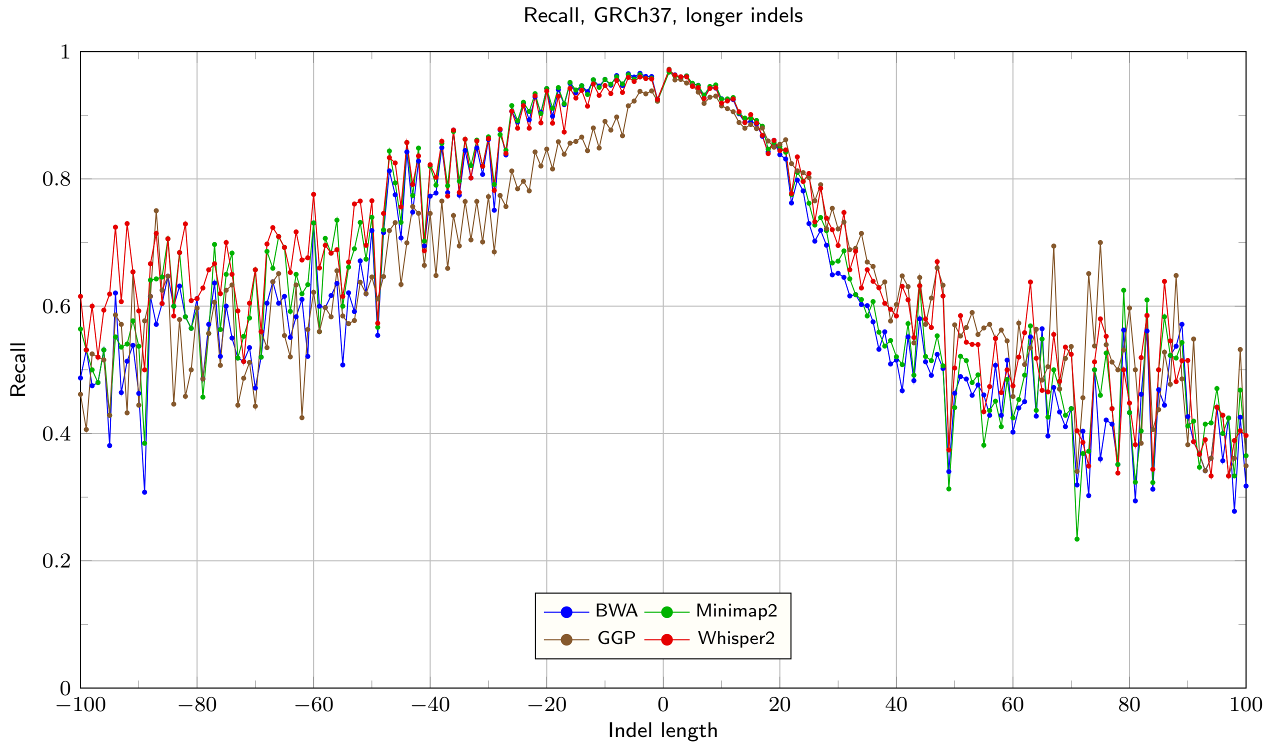

Figure 9: Recall for GRCh37, Strelka max. indel length set to 100

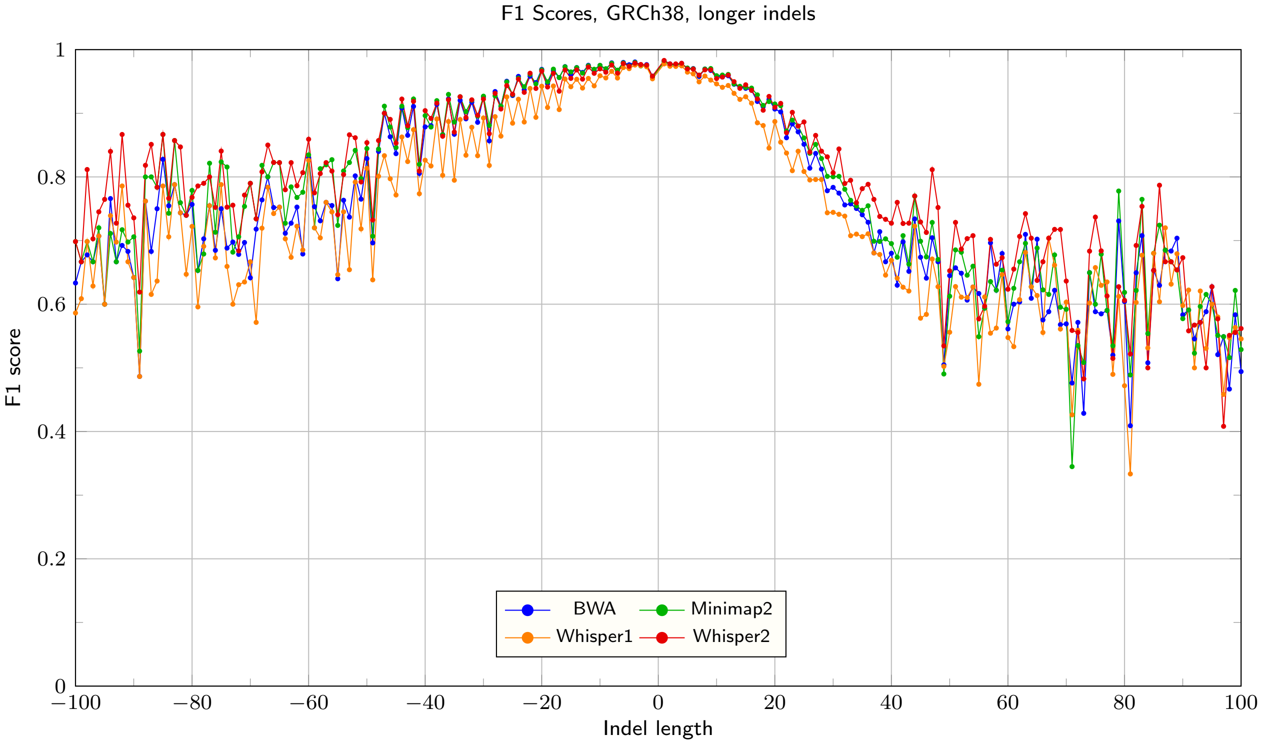

Figure 10: F1 Scores for GRCh38, Strelka max. indel length set to 100

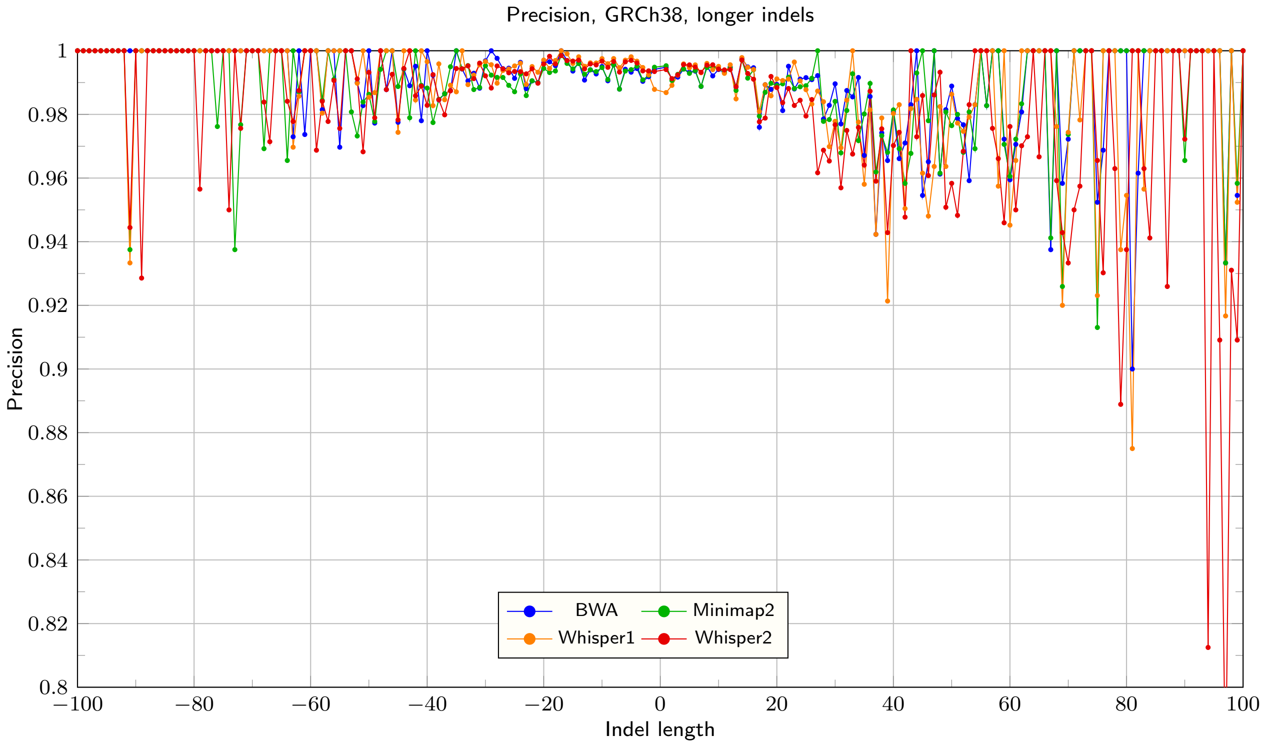

Figure 11: Precision for GRCh38, Strelka max. indel length set to 100

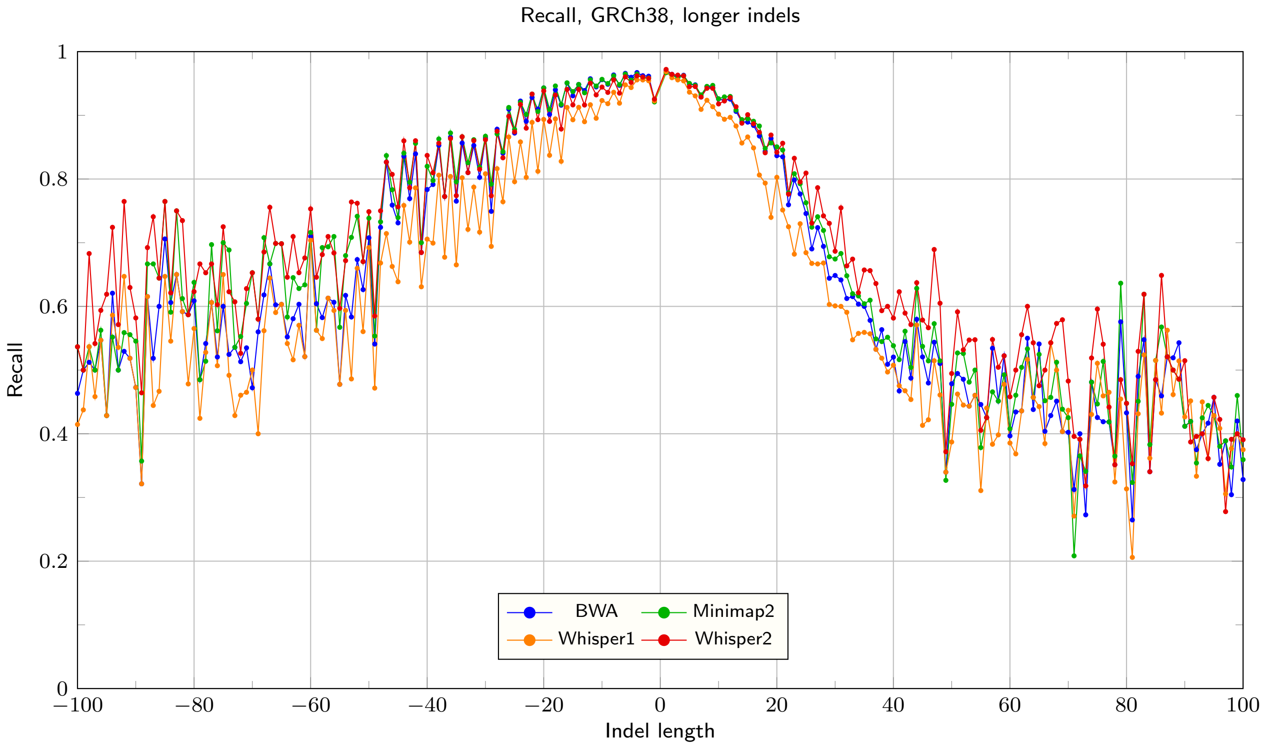

Figure 12: Recall for GRCh38, Strelka max. indel length set to 100

#### 5.3 SynDip evaluation for extended maximal indel length in Strelka, long indels

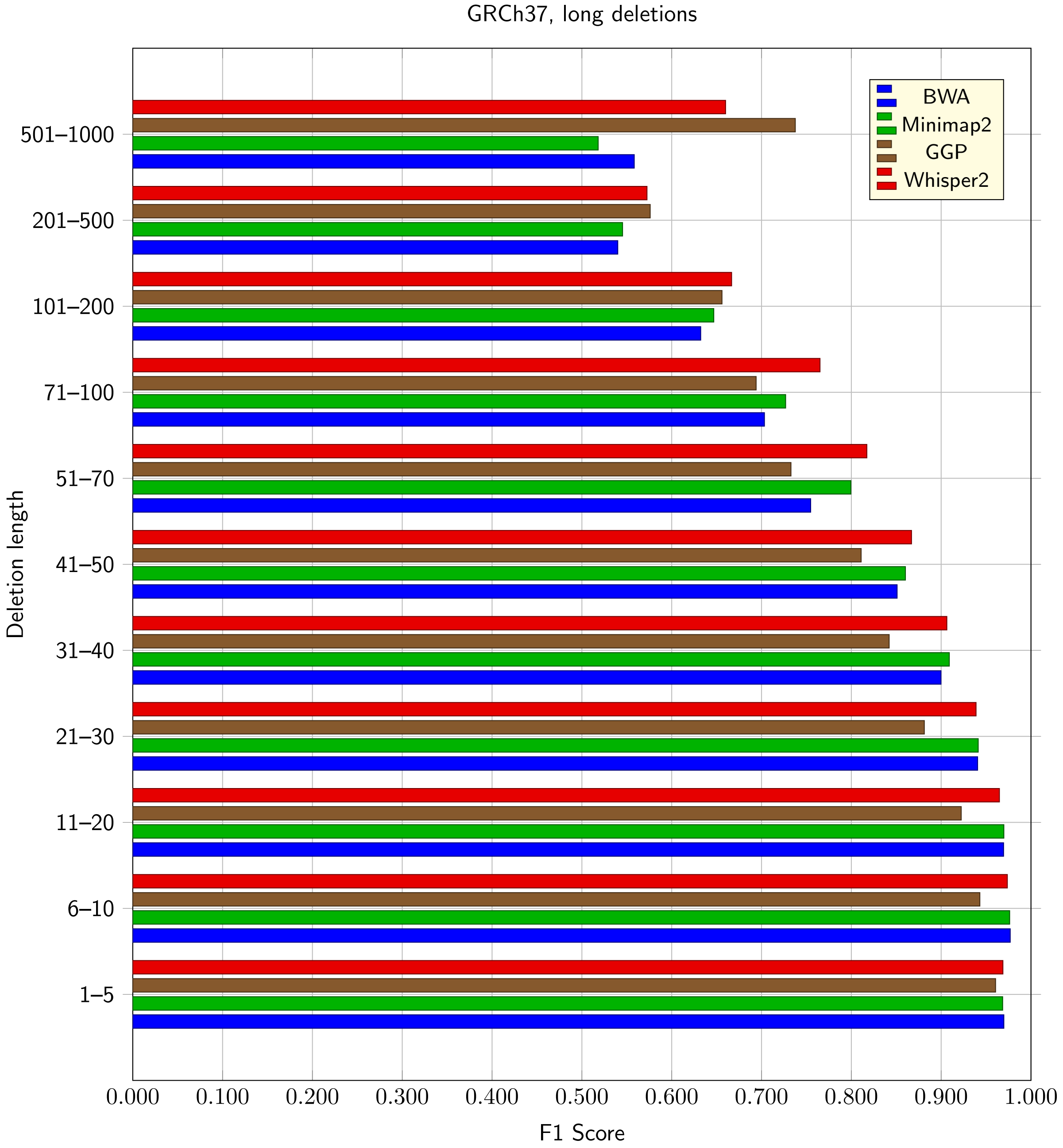

Figure 13: F1 Scores for deletions (in ranges) for GRCh37, Strelka max. indel length set to 100

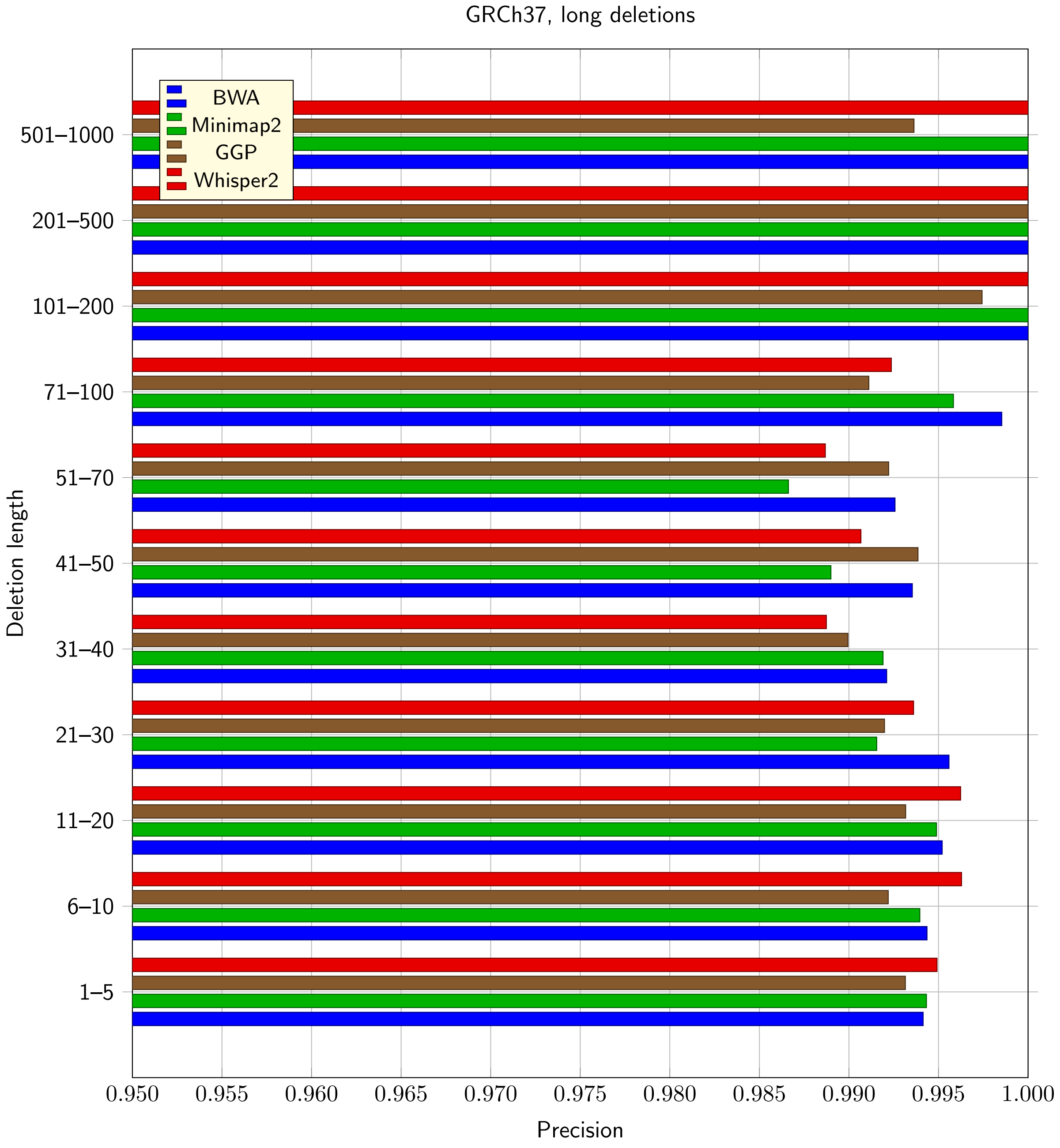

Figure 14: Precision for deletions (in ranges) for GRCh37, Strelka max. indel length set to 100

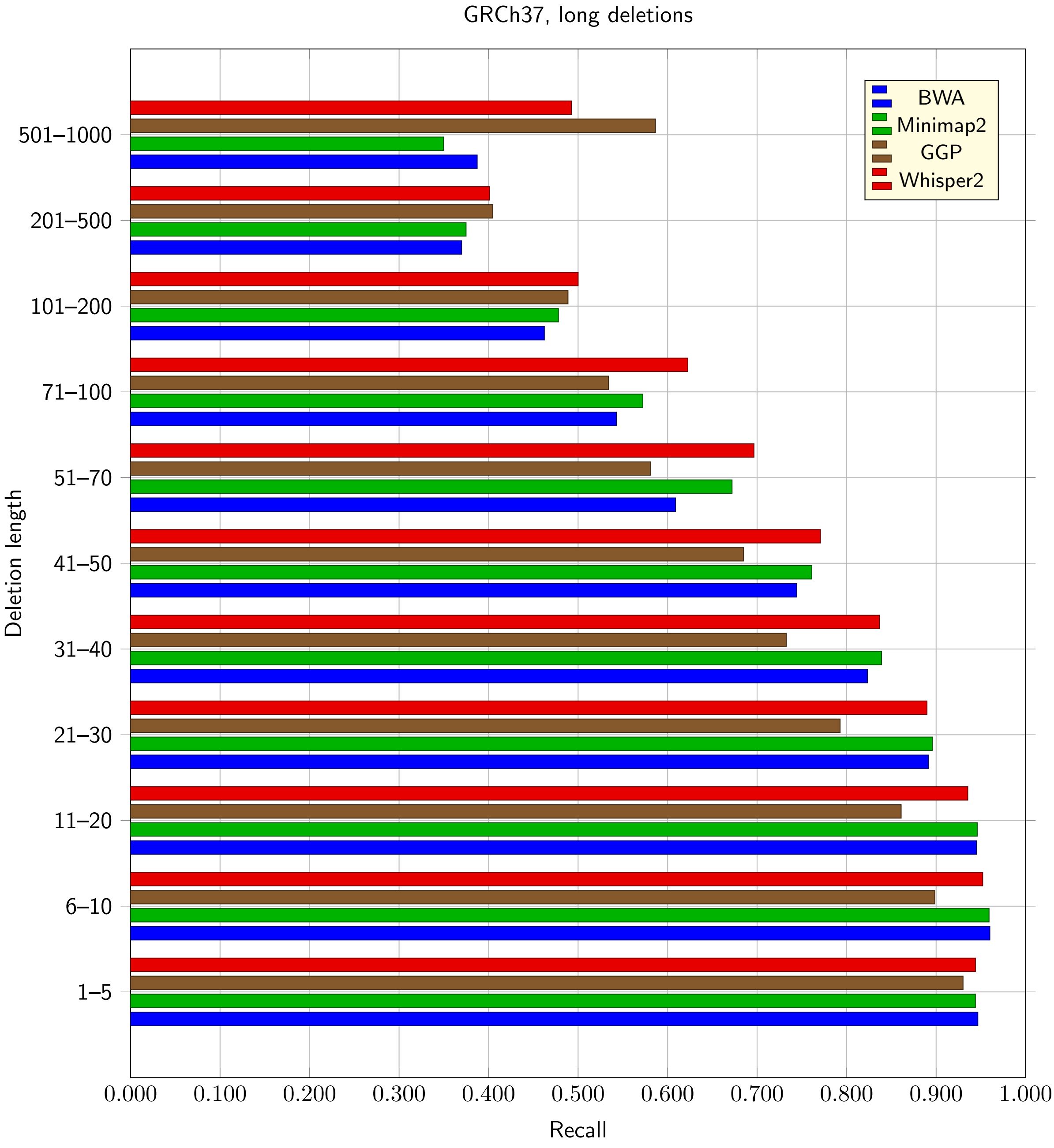

Figure 15: Recall for deletions (in ranges) for GRCh37, Strelka max. indel length set to 100

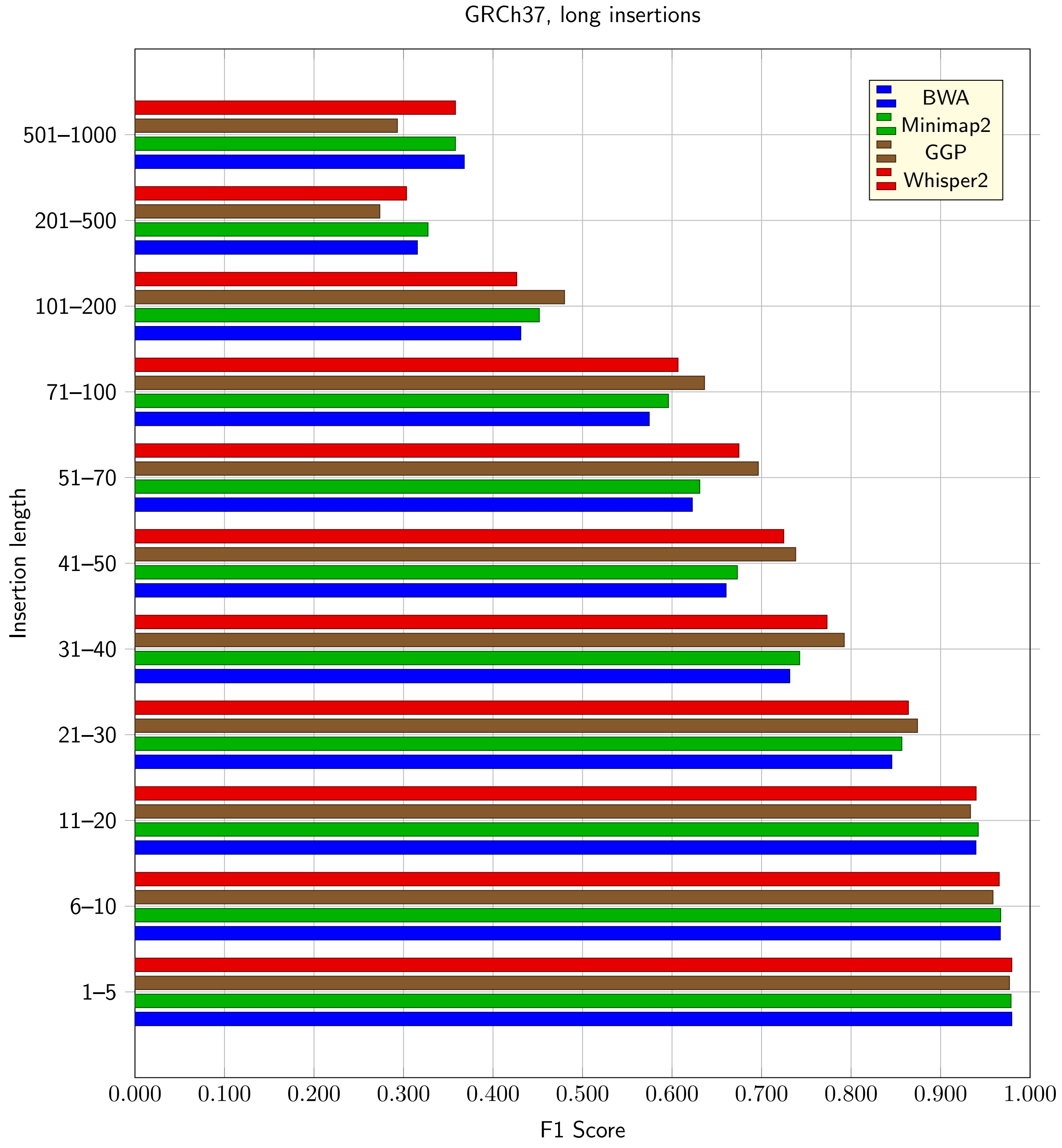

Figure 16: F1 Scores for insertions (in ranges) for GRCh37, Strelka max. indel length set to 100

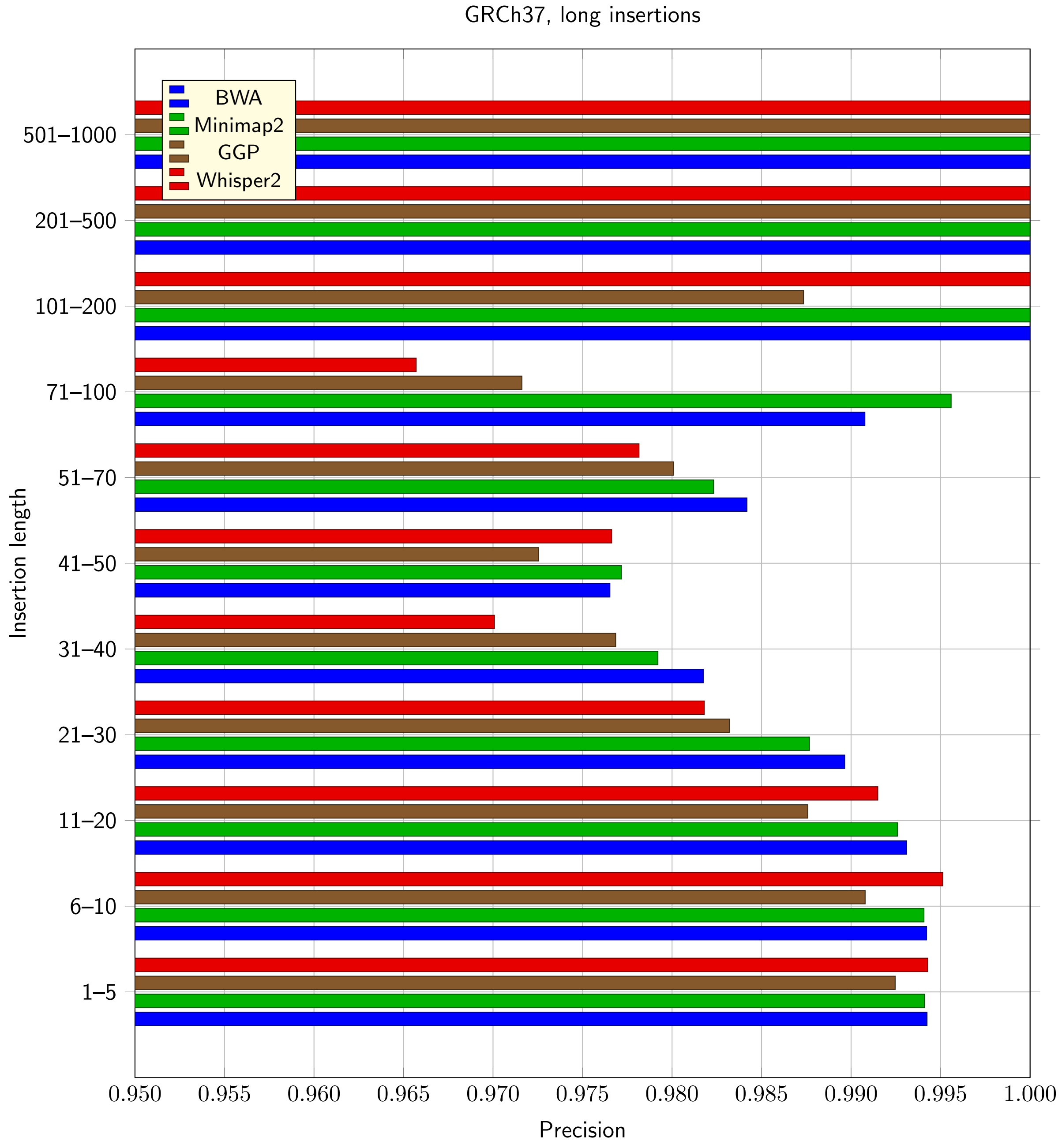

Figure 17: Precision for insertions (in ranges) for GRCh37, Strelka max. indel length set to 100

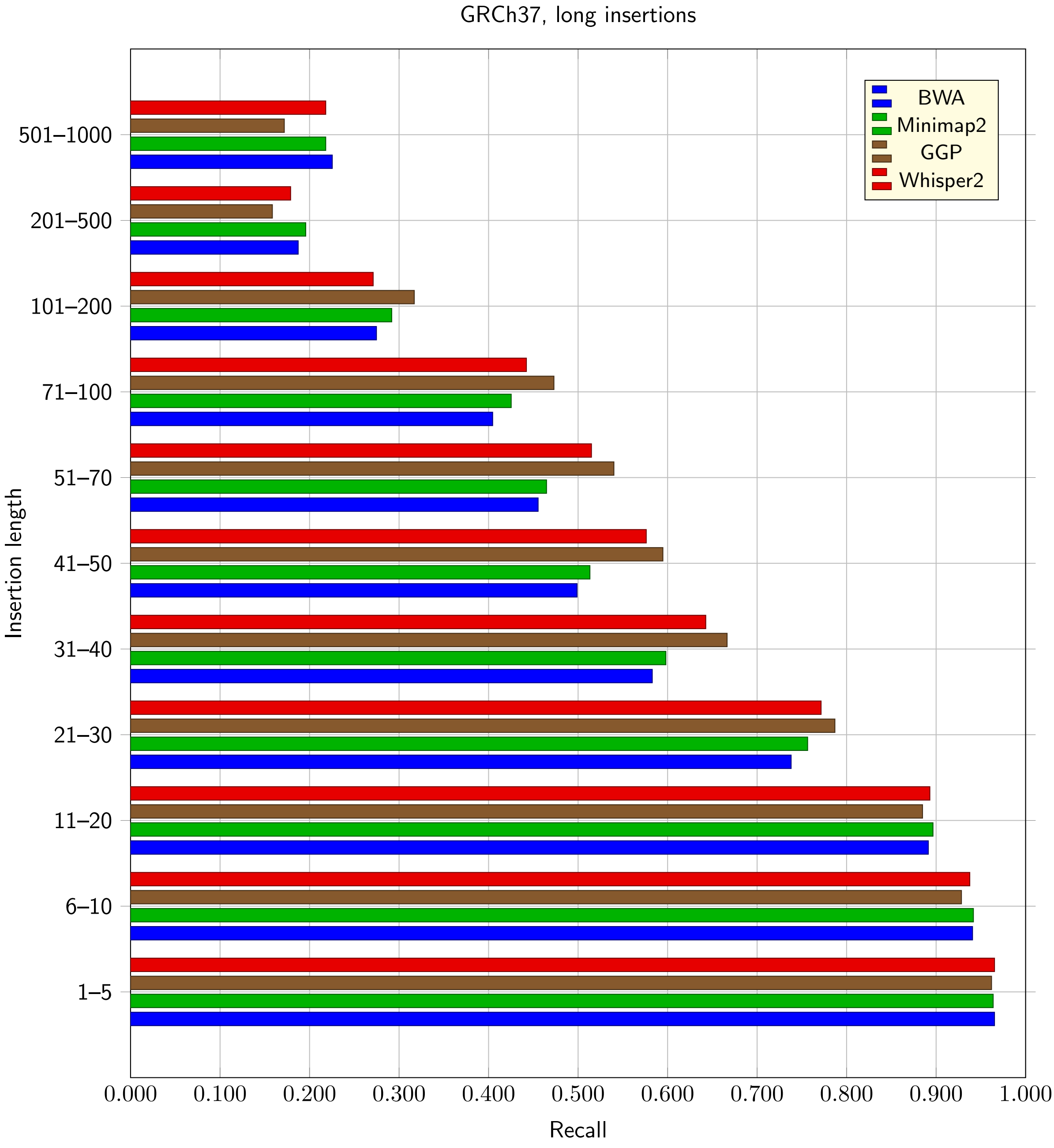

Figure 18: Recall for insertions (in ranges) for GRCh37, Strelka max. indel length set to 100

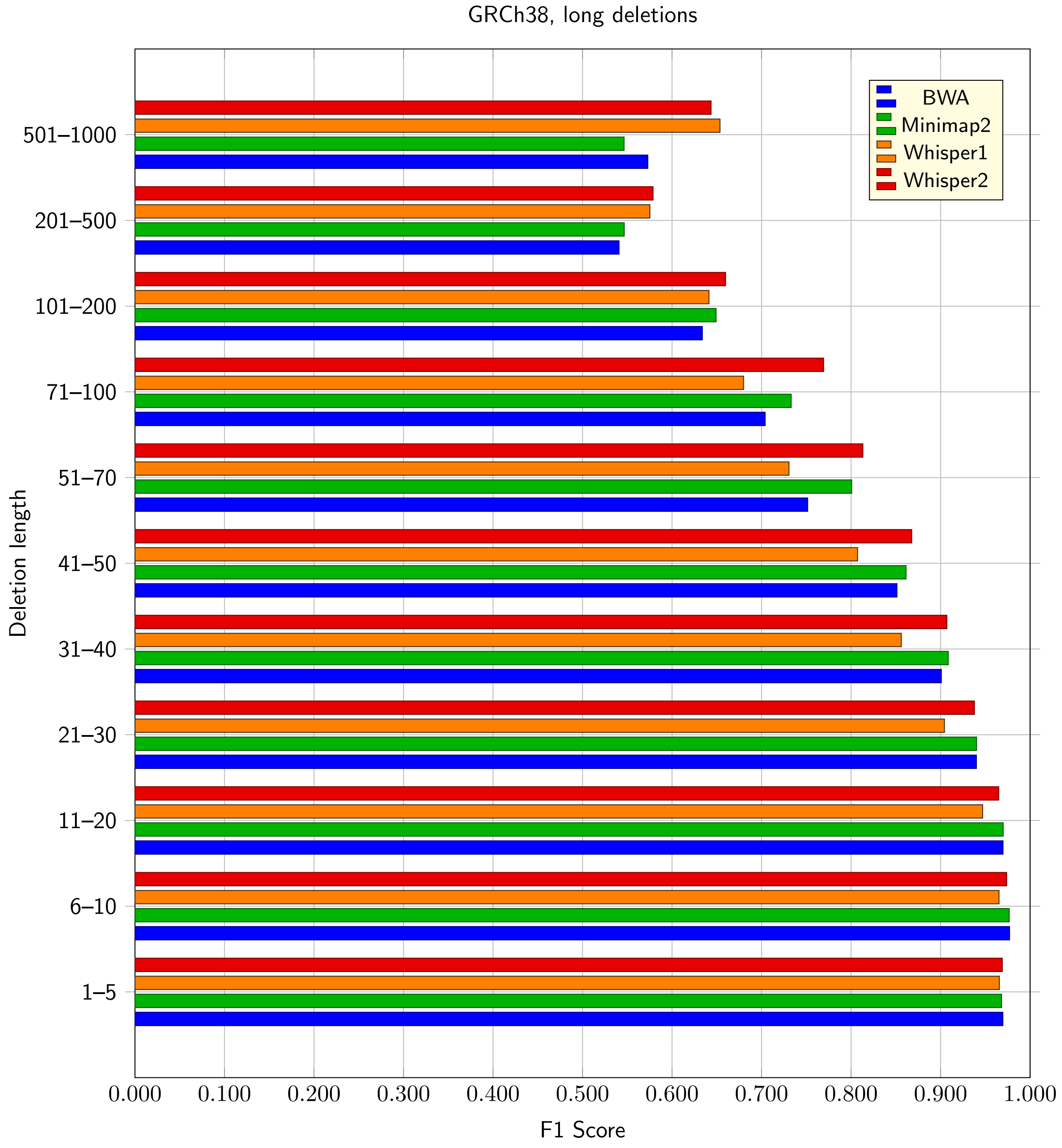

Figure 19: F1 Scores for deletions (in ranges) for GRCh38, Strelka max. indel length set to 100

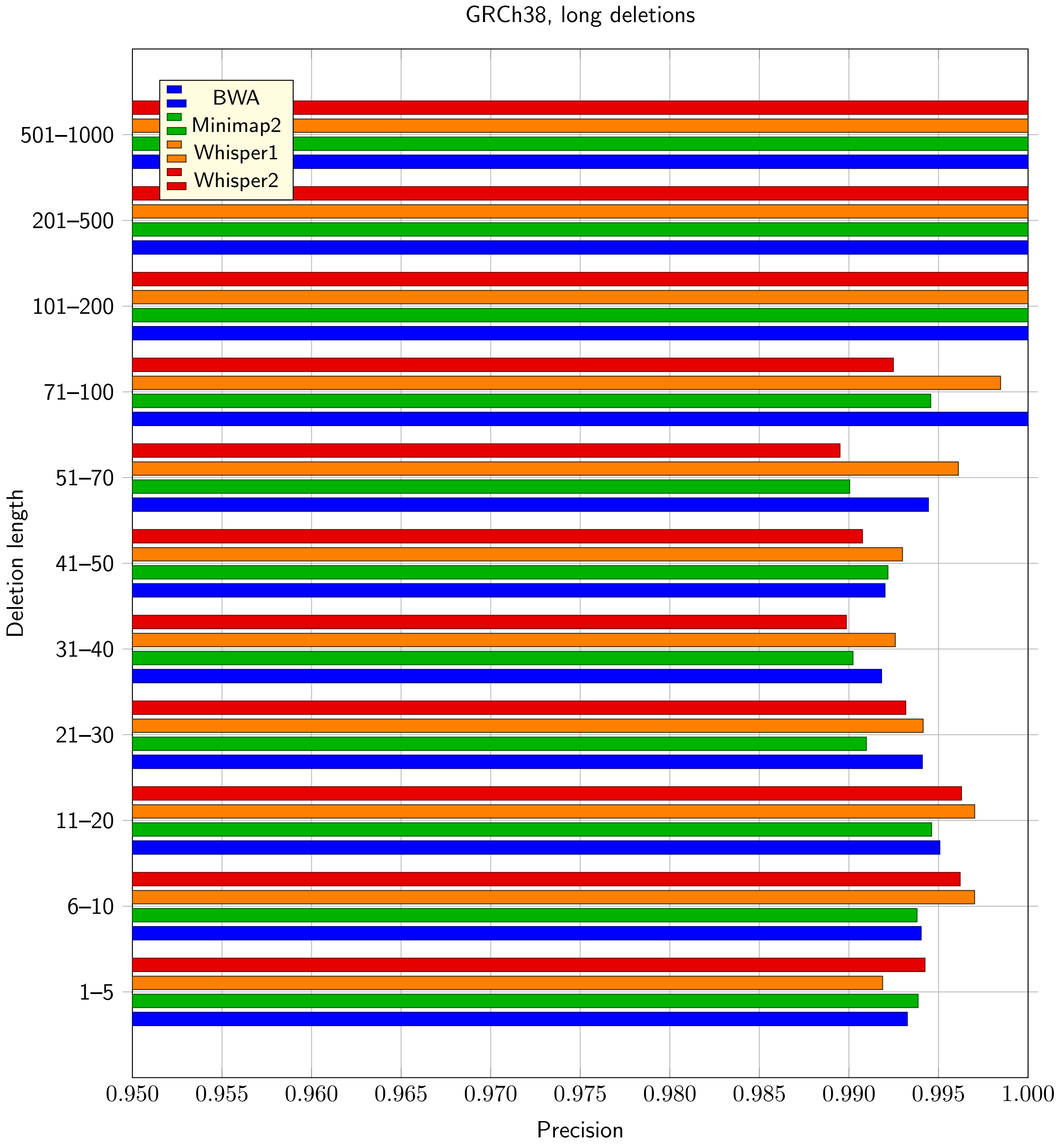

Figure 20: Precision for deletions (in ranges) for GRCh38, Strelka max. indel length set to 100

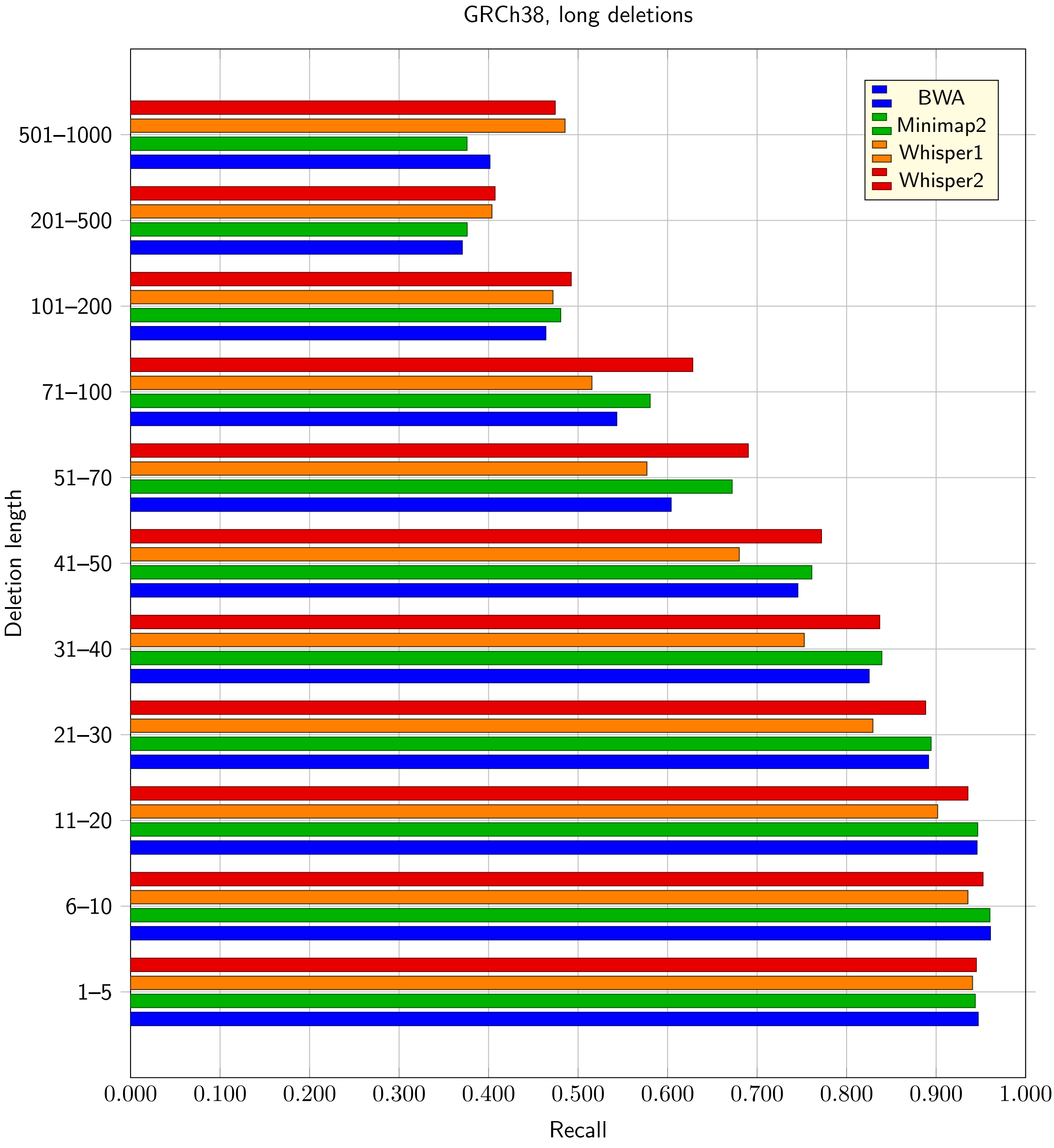

Figure 21: Recall for deletions (in ranges) for GRCh38, Strelka max. indel length set to 100

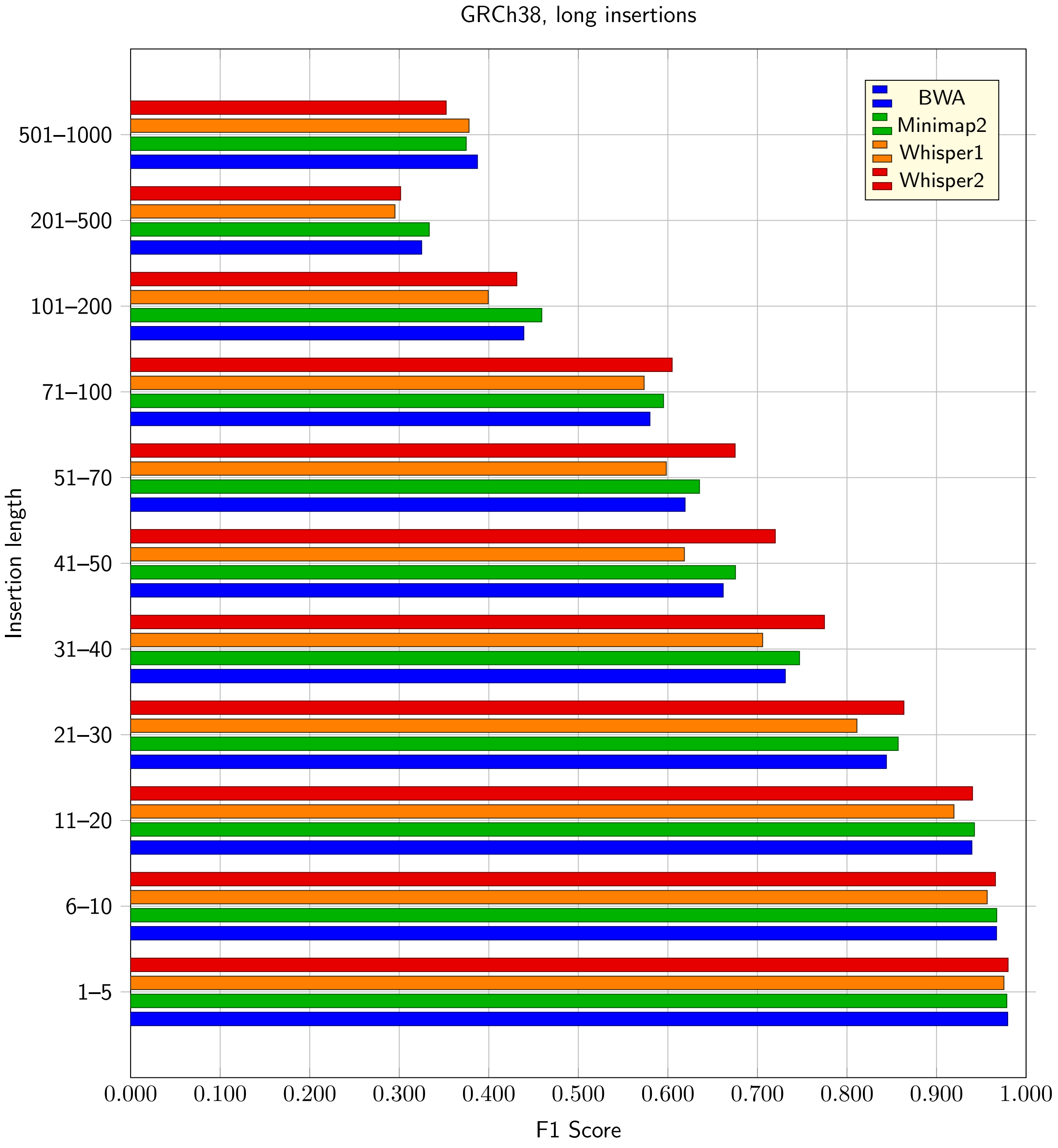

Figure 22: F1 Scores for insertions (in ranges) for GRCh38, Strelka max. indel length set to 100

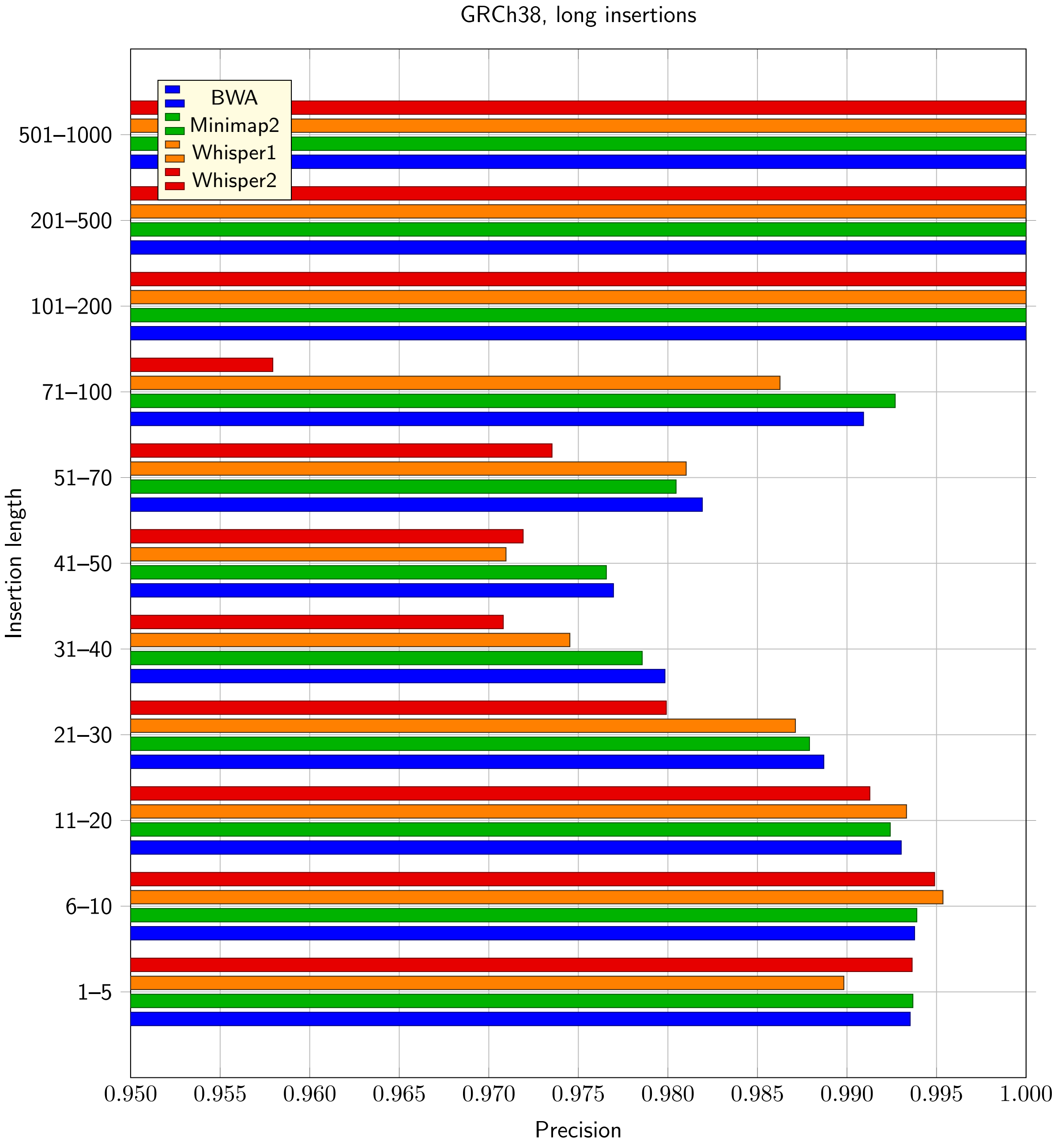

Figure 23: Precision for insertions (in ranges) for GRCh38, Strelka max. indel length set to 100

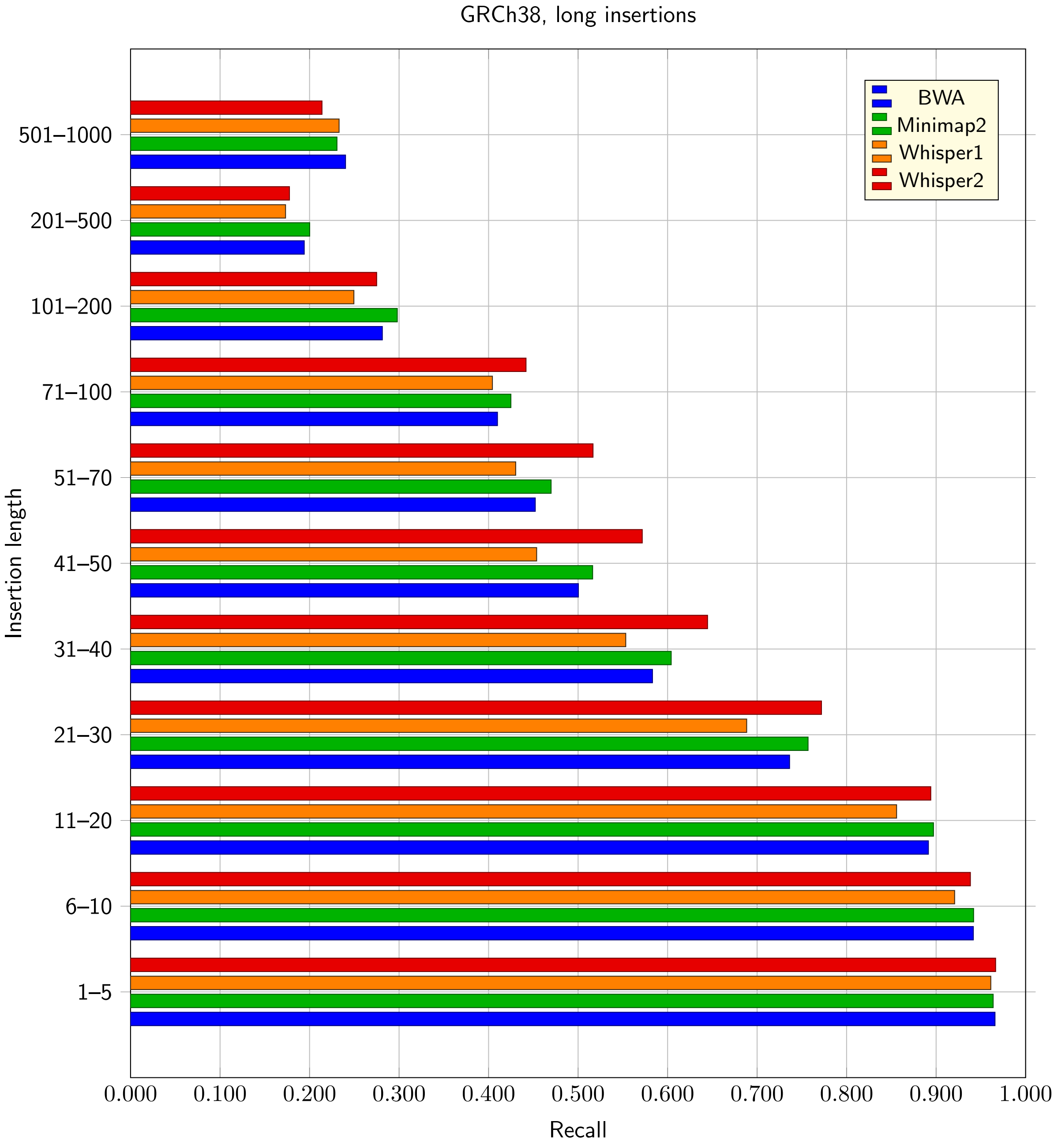

Figure 24: Recall for insertions (in ranges) for GRCh38, Strelka max. indel length set to 100

#### 5.4 Comparison of F1 Scores for default and extended maximal indel size in Strelka

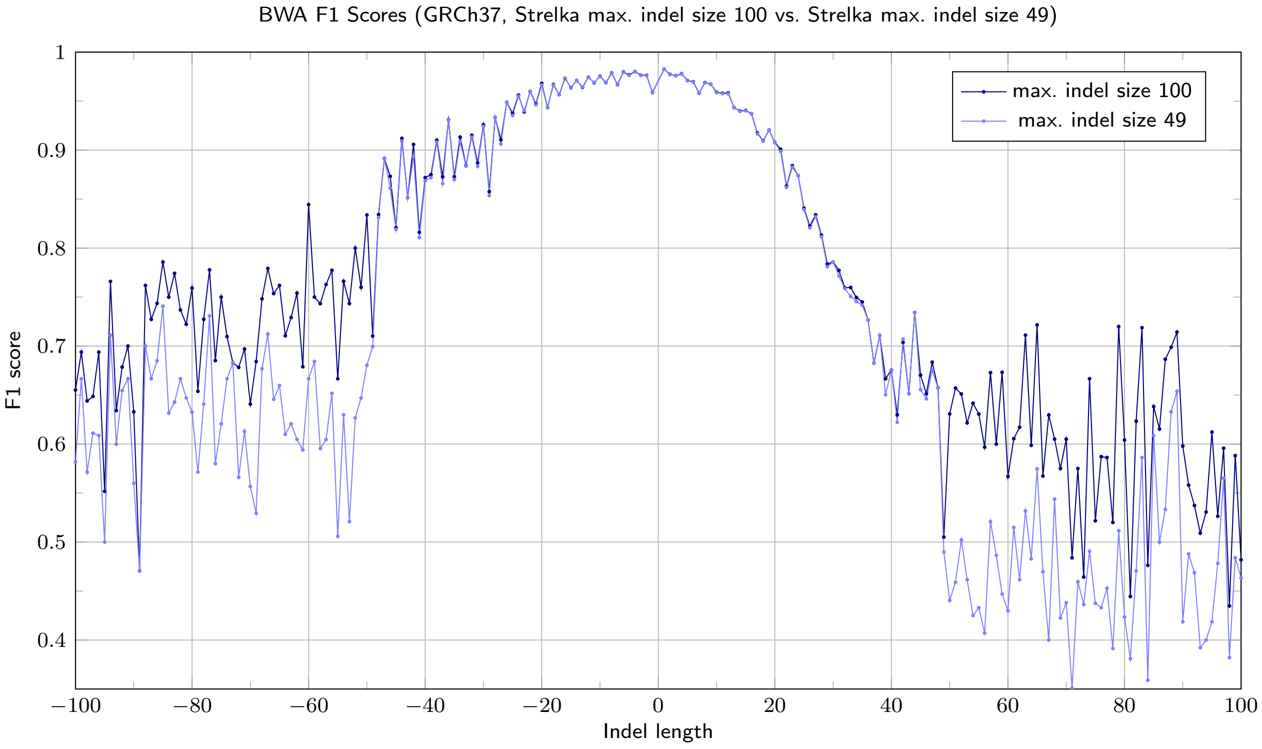

Figure 25: Comparison of BWA F1 Scores for default and Strelka max. indel length set to 100, GRCh38

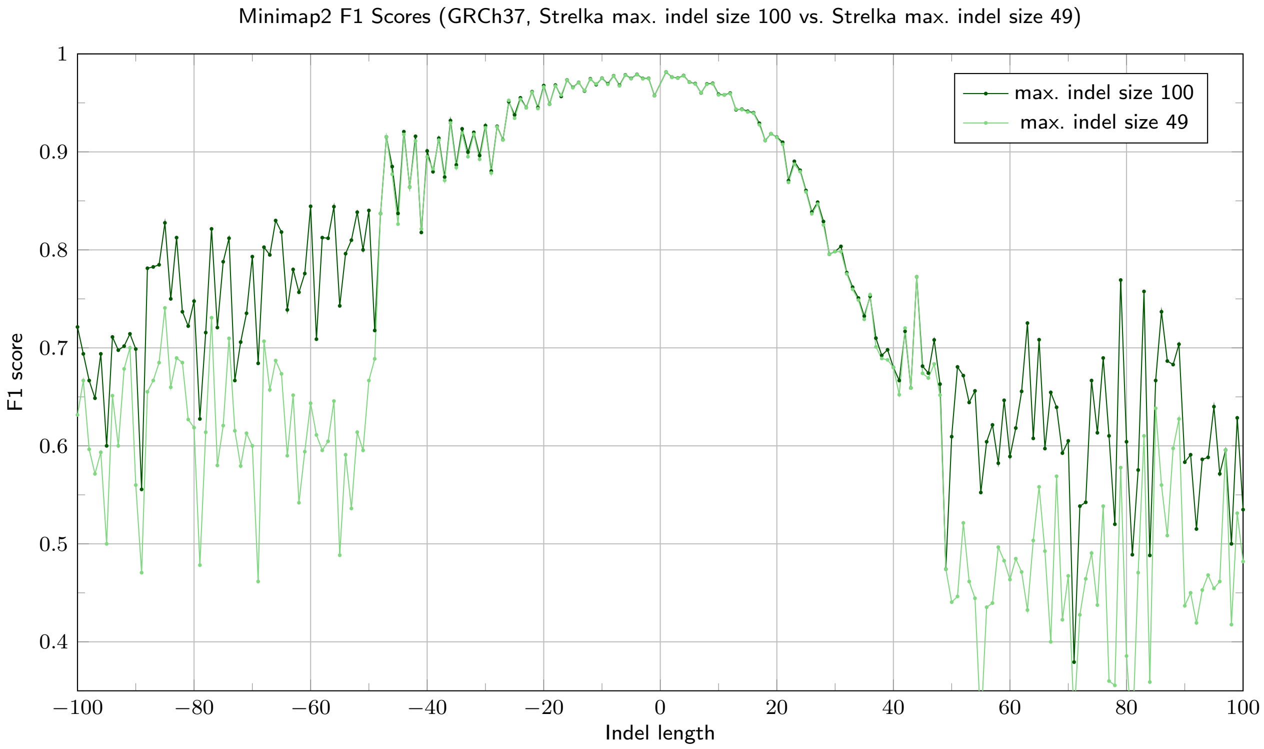

Figure 26: Comparison of Minimap F1 Scores for default and Strelka max. indel length set to 100, GRCh38

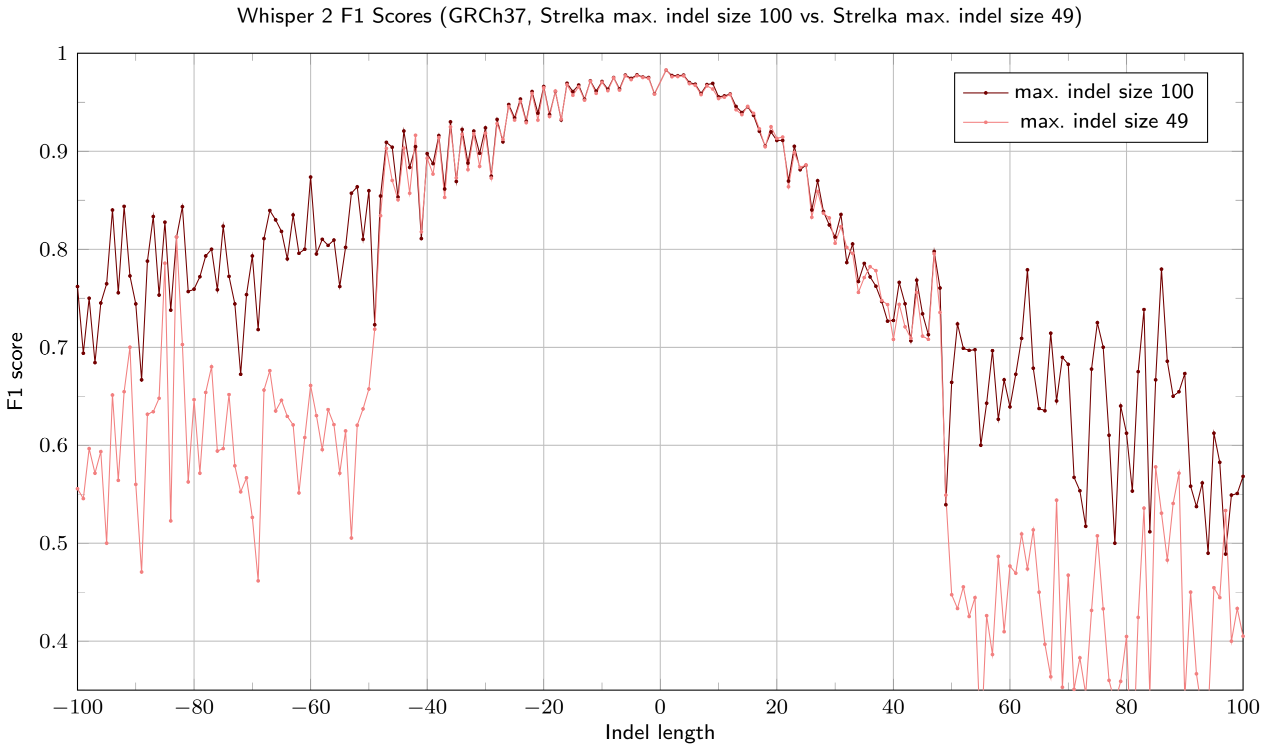

Figure 27: Comparison of Whisper 2 F1 Scores for default and Strelka max. indel length set to 100, GRCh38

#### 5.5 Comparison of advantages of the mappers over BWA

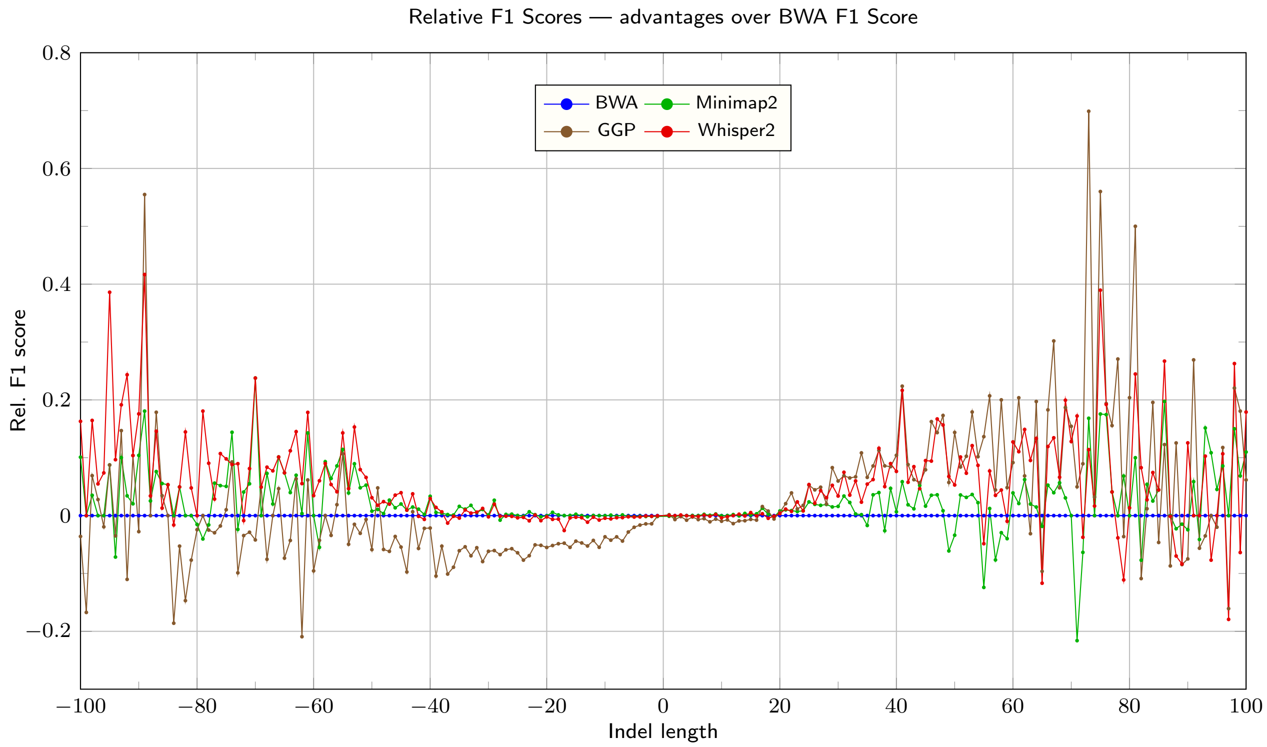

Figure 28: Relative advantage (mapper F1 Score divided by F1 Score minus 1) of the mappers over BWA — Strelka max. indel length set to 100, GRCh38
